# A chromosome-level assembly of the cat flea genome uncovers rampant gene duplication and genome size plasticity

**DOI:** 10.1101/2020.04.14.038018

**Authors:** Timothy P. Driscoll, Victoria I. Verhoeve, Joseph J. Gillespie, J. Spencer Johnston, Mark L. Guillotte, Kristen E. Rennoll-Bankert, M. Sayeedur Rahman, Darren Hagen, Christine G. Elsik, Kevin R. Macaluso, Abdu F. Azad

## Abstract

**Background:** Fleas (Insecta: Siphonaptera) are small flightless parasites of birds and mammals; their blood-feeding can transmit many serious pathogens (i.e. the etiological agents of bubonic plague, endemic and murine typhus). The lack of flea genome assemblies has hindered research, especially comparisons to other disease vectors. Accordingly, we sequenced the genome of the cat flea, *Ctenocephalides felis*, an insect with substantial human health and veterinary importance across the globe.

**Results:** By combining Illumina and PacBio sequencing with Hi-C scaffolding techniques, we generated a chromosome-level genome assembly for *C*. *felis*. Unexpectedly, our assembly revealed extensive gene duplication across the entire genome, exemplified by ∼38% of protein-coding genes with two or more copies and over 4,000 tRNA genes. A broad range of genome size determinations (433-551 Mb) for individual fleas sampled across different populations supports the widespread presence of fluctuating copy number variation (CNV) in *C. felis*. Similarly broad genome sizes were also calculated for individuals of *Xenopsylla cheopis* (Oriental rat flea), indicating that this remarkable “genome-in-flux” phenomenon could be a siphonapteran-wide trait. Finally, from the *C. felis* sequence reads we also generated closed genomes for two novel strains of *Wolbachia*, one parasitic and one symbiotic, found to co-infect individual fleas.

**Conclusion:** Rampant CNV in *C*. *felis* has dire implications for gene-targeting pest control measures and stands to complicate standard normalization procedures utilized in comparative transcriptomics analysis. Coupled with co-infection by novel *Wolbachia* endosymbionts – potential tools for blocking pathogen transmission – these oddities highlight a unique and underappreciated disease vector.

## Background

With over 2,500 described species, fleas (Hexapoda: Siphonaptera) are small (∼3 mm) flightless insects that parasitize mainly mammals and birds [1]. Diverging from Order Mecoptera (scorpionflies and hangingflies) in the Jurassic period [2], fleas are one of 11 extant orders of Holometabola, a superorder of insects that collectively go through distinctive larval, pupal, and adult stages. The limbless, worm-like flea larvae contain chewing mouthparts and feed primarily on organic debris, while adult mouthparts are modified for piercing skin and sucking blood. Other adaptations to an ectoparasitic lifestyle include wing loss, extremely powerful hind legs for jumping, strong claws for grasping, and a flattened body that facilitates movement on host fur and feathers.

The Oriental rat flea, *Xenopsylla cheopis*, and to a lesser extent the cat flea, *Ctenocephalides felis*, transmit *Yersinia pestis*, the causative agent of bubonic plague [3–5]. Fleas that feed away from their primary hosts (black rats and other murids) can introduce *Y*. *pestis* to humans, which historically has eliminated a substantial fraction of the world’s human population; e.g., the Plague of Justinian and the Black Death [5]. Bubonic plague remains a significant threat to human health [6, 7] as do other noteworthy diseases propagated by flea infestations, including murine typhus (*Rickettsia typhi*), murine typhus-like illness (*R*. *felis*), cat-scratch disease (*Bartonella henselae*), and Myxomatosis (myxoma virus) [8, 9]. Fleas also serve as intermediate hosts for certain medically-relevant helminths and trypanosome protozoans [10]. In addition to the potential for infectious disease transmission, flea bites are also a significant nuisance and can lead to serious dermatitis for both humans and their companion animals. Epidermal burrowing by the jigger flea, *Tunga penetrans*, causes a severe inflammatory skin disease known as Tungiasis, which is a scourge on many human populations within tropical parts of Africa, the Caribbean, Central and South America, and India [11, 12]. Skin lesions that arise from flea infestations also serve as sites for secondary infection. Collectively, fleas inflict a multifaceted human health burden with enormous public health relevance [13].

Most flea species reproduce solely on their host; however, their ability to feed on a range of different animals poses a significant risk for humans cohabitating with pets that are vulnerable to flea feeding – which includes most warm-blooded, hairy vertebrates [14]. As such, fleas also have a substantial economic impact from a veterinary perspective [15]. Many common pets are susceptible to flea infestations that often cause intense itching, bleeding, hair loss, and potential development of flea allergy dermatitis, an eczematous itchy skin disease. In the United States alone, annual costs for flea-related veterinary bills tally approximately $4.4 billion, with another $5 billion for prescription flea treatment and pest control [16]. Despite intense efforts to control infestations, fleas continue to pose a significant burden to companion animals and their owners [17].

Notwithstanding their tremendous impact on global health and economy, fleas are relatively understudied compared to other arthropod disease vectors [18]. While transcriptomics data for mecopteroids (Mecoptera + Siphonaptera) have proven useful for Holometabola phylogeny estimation [2], assessment of flea immune pathways [19], and analysis of opsin evolution [20], the lack of mecopteroid genomes limits further insight into the evolution of Antliophora (mecopteroids + Diptera (true flies)) and severely restricts comparative studies of disease vectors. Thus, sequencing flea genomes stands to greatly improve our understanding of the shared and divergent mechanisms underpinning flea and fly vectors, a collective lineage comprised of the deadliest animals known to humans [21]. To address this glaring void in insect genomics and vector biology, we sequenced the genome of *C*. *felis*, a principal vector of *R*. *typhi*, *R*. *felis*, and *Bartonella* spp. [22–25] and an insect with substantial human health and veterinary importance across the globe [1]. To overcome the minute body size of individual fleas, we pooled multiple individuals to generate sufficient DNA for sequencing, sampled from an inbred colony to reduce allelic variation, and applied orthogonal informatics approaches to account for challenges arising from the potential misassembly of haplotypes.

## Results

Pooled female fleas from the Elward Laboratory colony (Soquel, California; hereafter EL fleas) were used to generate short (Illumina), long (PacBio), and chromatin-linked (Hi-C) sequencing reads. A total of 7.2 million initial PacBio reads were assembled into 16,622 contigs (773.8 Mb; N50 = 61 Kb), polished with short-read data, then scaffolded using Hi-C into 3,926 scaffolds with a final N50 of 71.7 Mb. A total of 193 scaffolds were identified as arising from microbial sources and removed before gene model prediction and annotation. A large fraction of the total assembly (85.6%, or 654 Mb) was found in nine scaffolds (all greater than 10 Mb, hereafter BIG9), while the remaining 14.4% (119.8 Mb) comprised scaffolds less than 1 Mb in length; therefore, we suggest the *C*. *felis* genome contains nine chromosomes (**Fig. 1A**), an estimate consistent with previously determined flea karyotypes [26, 27]. The 3,724 shorter scaffolds (all less than 1 Mb) mapped back to unique locations on BIG9 scaffolds (**Additional file 1: Fig. S1A**) but were not assembled into the BIG9 scaffolds via proximity ligation. Comparison of *C*. *felis* protein-encoding genes to the Benchmarking Universal Single-Copy Orthologues (BUSCO [28]) for eukaryotes, arthropods, and insects indicates our BIG9 assembly is robust and lacks only a few conserved genes (**Additional file 1: Fig. S1B**). As a result, we focus our subsequent analyses on the BIG9 scaffolds unless otherwise noted.

**Fig. 1.**
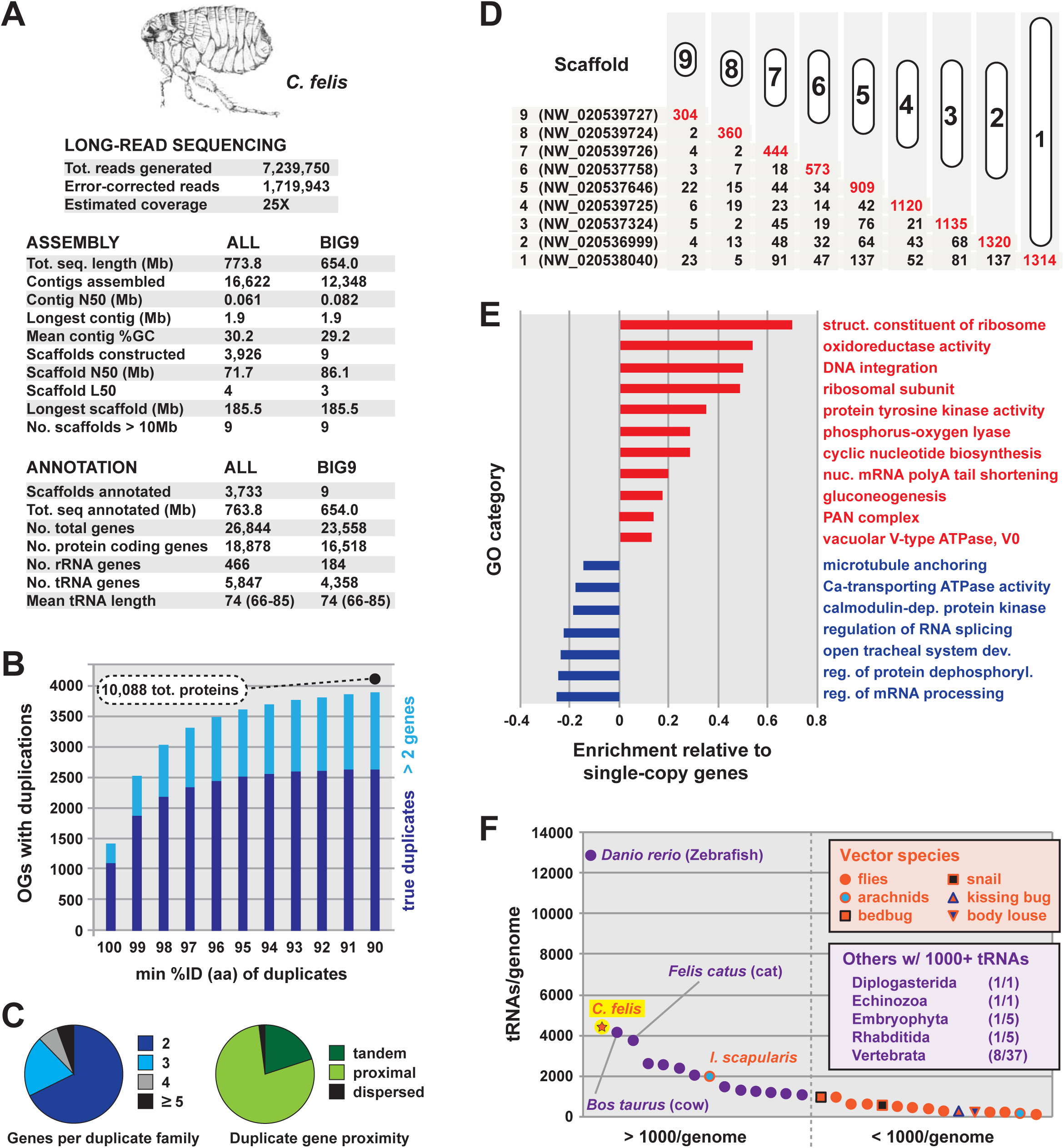
*C*. *felis* genome characteristics. (**A**) Summary statistics for long-read sequencing, assembly and gene annotation. (**B**) Of 16,518 total protein-encoding genes (BIG9 scaffolds), 10,088 are derived from gene duplications (6,225 duplication events within 3,863 OGs at a threshold of 90% aa identity). (**C**) Assessment of the number of genes per duplication (*left*) and the relative distances between duplicate genes (*right*). Distances were computed only for true duplications (n=2 genes) at a threshold of 90% aa identity. (**D**) Gene duplications are enriched within BIG9 scaffolds (tandem and proximal, red numbers) versus across scaffolds (dispersed, black numbers). (**E**) Enriched cellular functions of duplicate genes relative to single-copy genes. (**F**) *C*. *felis* belongs to a minimal fraction of eukaryotes containing abundant tRNA genes. tRNA gene counts are shown for disease vectors (VectorBase [84]) and eukaryotes carrying over 1000 tRNA genes (GtRNAdb [29]; ratios show number of genomes with > 1000 tRNA genes per taxon.

### The *C*. *felis* genome and unprecedented gene duplication

Previous work using flow cytometry estimated the size of the female *C. felis* genome at 465 Mb, while our BIG9 assembly contained 654 Mb total bases (25% larger). Furthermore, BUSCO analysis suggested that roughly 30% of conserved, single-copy Insecta genes in the BUSCO set were duplicated in our assembly (**Additional file 1: Fig. S1B**). In order to investigate whether this duplication might be widespread across the genome, and thereby account for the larger size of our assembly, we used BLASTP to construct *C. felis*-specific protein families at varying levels of sequence identity from 85-100%. Remarkably, 61% (10,088) of all protein-encoding genes in *C. felis* arise from duplications at the 90% identity threshold or higher (**Fig. 1B**). Over 68% of these comprise true (n=2) duplications, most of which occur as tandem or proximal loci less than 12 genes apart (**Fig. 1C, Additional file 1: Fig. S1L**). We observed little change in either the total number of duplications or the distribution at thresholds below 90% identity; consequently, we define “duplications” here as sequences that are 90% identical or higher (see **Methods**).

Duplications are on-going and rapidly diverging as evinced by: 1) their high concentration on individual BIG9 scaffolds (**Fig. 1D**, **Additional file 1: Fig. S1C-K**), 2) a lack of increasing divergence with greater distance on scaffolds (**Additional file 1: Fig. S1L**), and 3) a lack of increasing divergence for duplicate genes found across different scaffolds (**Additional file 1: Fig. S1M**). Among cellular functions for duplicate genes, certain transposons and related factors (GO:0015074, “DNA integration”) are enriched relative to 6,430 single copy protein-encoding genes (**Fig. 1E**, **Additional file 2: Table S1**). However, the frequency and distribution of these elements are dwarfed by total duplicate genes (**Additional file 1: Fig. S1N**). Additionally, transposons and other repeat elements encompass only 10% of the genome (**Additional file 1: Fig. S1O**), indicating that selfish genetic elements do not contribute significantly to the rampant gene duplication observed. Thus, the *C*. *felis* genome is remarkable given that genes producing duplications (n=3,863 or ∼38% of total protein-encoding genes) are 1) indiscriminately dispersed across chromosomes, 2) not clustered into blocks that would suggest whole or partial genome duplications, and 3) not the product of repeat element-induced genome obesity.

The *C*. *felis* genome also carries an impressive number of tRNA-encoding genes (*n*=4,358 on BIG9 scaffolds) (**Fig. 1A**). While tRNA gene numbers and family compositions vary tremendously across eukaryotes [29], the occurrence of more than 1000 tRNA genes per genome is rare (**Fig. 1F**). Notably, the elevated abundance of tRNA genes in *C. felis* is complemented by an enrichment in translation-related functions among duplicated protein-coding genes (**Fig. 1E, Additional file 2: Table S1**). While this possibly indicates increased translational requirements to accommodate excessive gene duplication, it is more likely a consequence of the indiscriminate nature of the gene duplication process. Relative to tRNA gene frequencies in other holometabolan genomes, *C*. *felis* has several elevated (Arg, Val, Phe, Thr) and reduced (Gly, Pro, Asp, Gln) numbers of tRNA families (**Additional file 1: Fig. S1P**); however, *C*. *felis* codon usage is typical of holometabolan genomes (**Additional file 1: Fig. S1Q**). Like proliferated protein-encoding genes, the significance of such high tRNA gene numbers is unclear but further accentuates a genome in flux.

### Genome size estimation

Duplicated regions (including intergenic sequences) account for approximately 227 Mb of the *C. felis* genome; when subtracted from the BIG9 assembly (654 Mb), the resulting “core” genome size of 427 Mb is congruous with a previous flow cytometry-based genome size estimate (mean of 465 Mb, range of 32 Mb) for cat fleas previously assayed from a different geographic locale [30]. To determine if EL fleas possess a greater genome size due to pronounced gene duplication relative to other cat fleas, we similarly used flow cytometry to estimate genome sizes for individual EL fleas and compared them to the previous findings. As expected, mean genome size was not significantly different between sex-matched *C. felis* from the two populations (p = 0.1299). Remarkably, however, no two individual EL fleas possessed comparable genome sizes, with an overall uniform size distribution and relatively large variability (118 Mb) (**Fig. 2A**; **Additional file 3: Fig. S2**). Indeed, the coefficient of variation for *C. felis* (0.13; n = 26) was 3.2X higher than that of either *Drosophila melanogaster* (0.040; n = 26) or *D. viridis* (0.039; n = 26), which were prepared and measured concurrently (**Fig. 2A**, inset), underscoring the extraordinary extent of inter-individual variation in *C. felis*. Genome size estimates for another flea (the rat flea, *X*. *cheopis*, also sex-matched) show a similar uniform distribution and range across individuals (**Fig. 2A**), pointing to an extraordinary genetic mechanism that may define siphonapteran genomes.

**Fig. 2.**
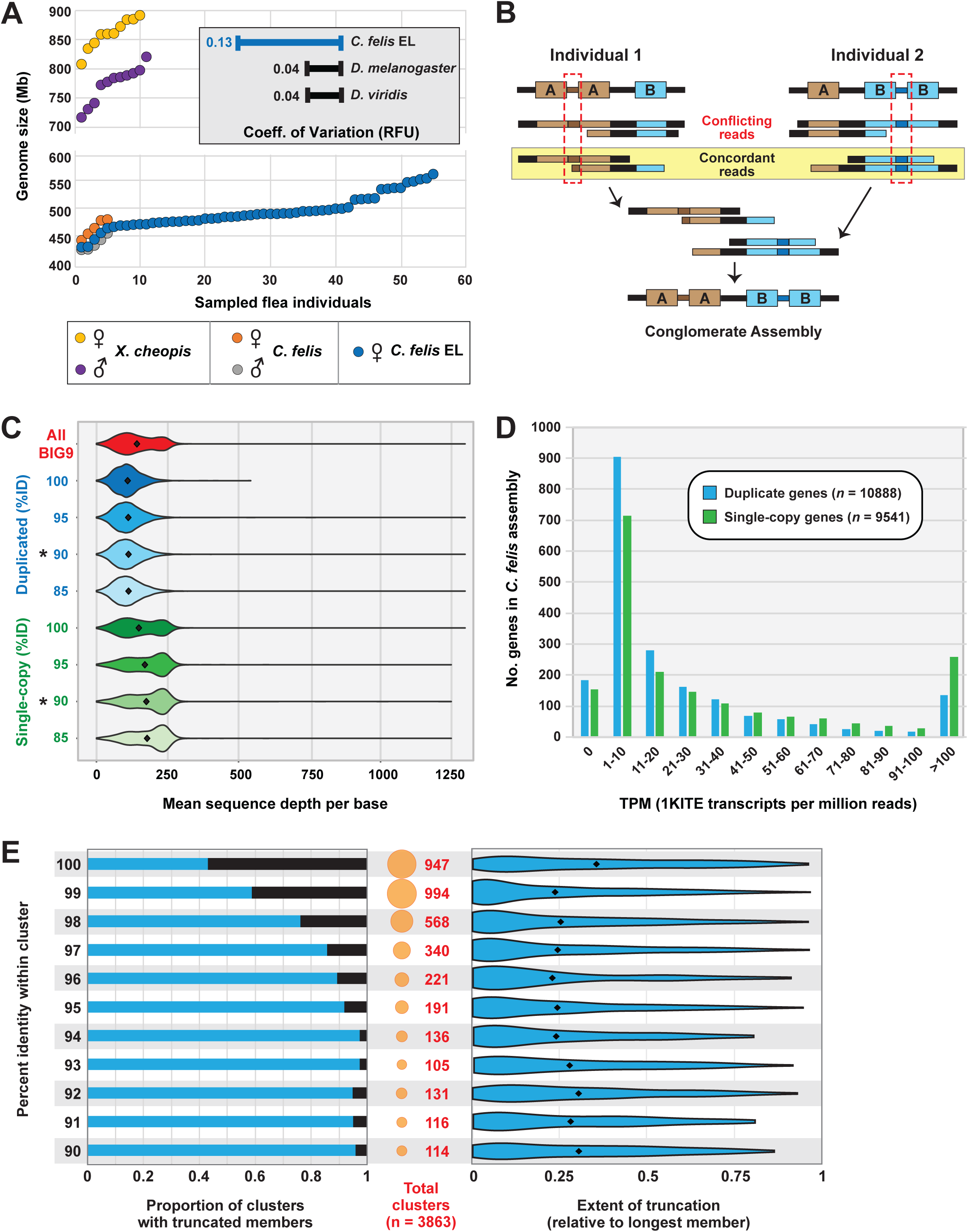
Evidence for excessive copy number variation in the *C*. *felis* genome. (**A**) Flea genome size estimates. Flow cytometer-based estimates were performed for male and female individuals of *X*. *cheopis* (Texas), *C*. *felis* (Texas), and for female *C. felis* EL from the sequenced colony (see **Additional file 3: Fig. S2**). The inset (top right) depicts the coefficients of variation in measured fluorescence (relative fluorescence units; RFU) for *Drosophila melanogaster* (n=26), *D. viridis* (n=26), and *C. felis* EL (n=26) females prepared and analyzed simultaneously. (**B**) Graphic depiction of assembling CNV. Two theoretical individual fleas are shown with different CNVs for loci A and B. Regions unique to each individual genome are shown by the red dashed boxes. Only reads concordant between individuals are included in the conglomerate assembly. (**C**) Comparison of Illumina read coverage-mapping between duplicate genes (blue) and single-copy genes (green) at different %ID thresholds. Reads that mapped to multiple locations (alternative mappings) were included. Asterisks indicate statistically significant difference (Welch Two-Sample t-test, p < 2.2e-16) between mean coverage of single-copy and duplicate genes at the 90 %ID threshold. (**D**) Transcriptional support for *C. felis* EL genes within the 1KITE transcriptomic data. Counts of transcripts per million reads (TPM) were mapped (Hisat2 & Stringtie), binned, and plotted against the number of duplicated (blue) and single-copy (green) genes in the BIG9 assembly. (**E**) Extent of truncation within clusters of duplicated genes in *C. felis*. The number of clusters with truncated members at each integer %ID threshold (left) was calculated as the proportion of total clusters at that threshold (center). The distribution of length differences in these clusters (relative to the longest member in each cluster) is plotted as a violin plot (right); black diamonds represent the mean length difference at each %ID threshold.

Accordingly, we propose that our assembly captured a conglomeration of individual flea copy number variations (CNV) that is cumulative for all expansions and contractions of duplicate regions (**Fig. 2B**). The presence of extensive gene duplications is further supported by mapping short read Illumina data to our assembly, which showed a significantly reduced mean read depth across duplicated loci versus single-copy genes (**Fig. 2C**). As an alternative to CNV, we considered that allelic variation could also be contributing to extensive gene duplication in our assembly. To address this concern, we took three approaches. First, polished contigs were scanned for haplotigs using the program *Purge Haplotigs* [31]; no allelic variants were detected. Second, we mapped 1KITE transcriptome reads [2] generated from fleas of an unrelated colony (Kansas State University) to our assembly (**Fig. 2D**). If our sequence duplication is a result of allelic variation within the EL colony, we would expect to see a lack of congruence in the distribution of transcripts mapping to single copy genes versus duplicates (different colonies with different allelic variation). We might also expect to see a significant proportion of transcripts that do not map at all. Instead, 91% of 1KITE reads map to CDS in our assembly, and the distributions of transcripts mapping to single copy and duplicate genes are identical.

Third, we reasoned if sequence duplications are the result of misassembled allelic variants, then most duplicate CDS within a cluster would be the same length. Alternatively, if duplications are true CNVs, we would expect a significant number of truncations as a consequence of gene purging associated with unequal crossing over. To assess this, we determined the proportion of duplicate clusters with one or more truncated members, as well as the extent of truncation relative to the longest member of the cluster (**Fig. 2E**). Approximately 70% of gene duplications are not comparable in length. In addition, mean extent of truncation is 25% or greater across all clusters regardless of % identity. Together with genome size estimations, short read mapping analysis, and transcript mapping to our assembly, these data indicate active gene expansion and contraction underpinning CNV in fleas and dispel allelic variation as a significant contributor to gene duplication. While the cytogenetic mechanisms are unclear, elevated numbers of DNA repair enzymes (GO:0006281) relative to genome size may correlate with excessive CNV (**Additional file 2: Table S1**).

### Genome evolution within Holometabola

Despite inordinate gene duplication, the completeness of the *C*. *felis* proteome as estimated by occurrence of 1,658 insect Benchmarking Universal Single-Copy Orthologues (BUSCOs) is congruous with those of other sequenced holometabolan genomes (**Fig. 3A**). Only one other genome (*Aedes albopictus*) contains greater gene duplication among BUSCOs than *C*. *felis*; however, this mosquito genome is much larger (∼2 Gb) and riddled with repeat elements [32]. A genome-wide analysis of shared orthologs among 53 holometabolan genomes indicates a slight affinity of *C*. *felis* with Coleoptera, though the divergent nature of Diptera and availability of only a single flea genome likely mask inclusion of fleas with flies (**Fig. 3B**). Overall, phylogenomics analysis reveals that *C*. *felis* harbors 3,491 orthologs found in at least one other taxon from each holometabolan order (**Fig. 3C**); however, only 577 “core” orthologs were present in all taxa from every order (**Fig. 3C**, yellow bar), reflecting either incomplete genome assemblies or an incredibly patchwork Holometabola accessory genome (**Additional file 4: Fig. S3A**). Other conserved protein-encoding genes that define higher-generic groups (**Fig. 3C**, inset) will inform lineage diversification within Holometabola (**Additional file 5: Table S2**). Conversely, 29 protein-encoding genes absent in *C*. *felis* but conserved in Panorpida species (Antliophora + Lepidoptera (butterflies and moths)) stand to illuminate patterns and processes of flea specialization via reduction (**Additional file 4: Fig. S3B**, **Additional file 5: Table S2**). Overall, despite its parasitic lifestyle and reductive morphology, *C*. *felis* has not experienced a significant reduction in gene families (**Additional file 4: Fig. S3A**, **Additional file 5: Table S2**) as seen in other host-dependent eukaryotes [33].

**Fig. 3.**
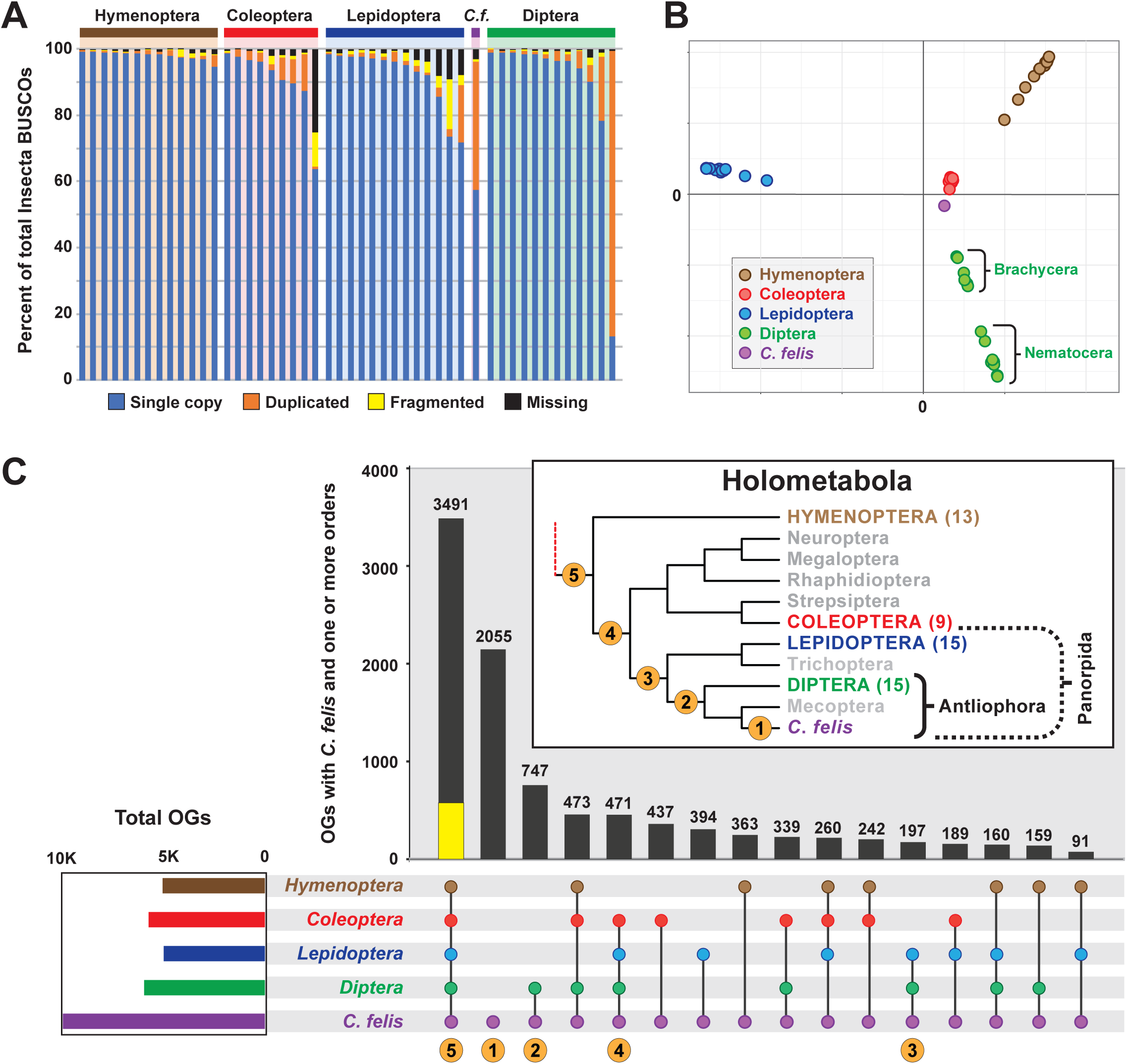
Phylogenomics analysis of the *C*. *felis* genome. (**A**) Assessing completeness and conservation of select holometabolan genomes using insect (n=1,658) Benchmarking Universal Single-Copy Orthologues (BUSCOs) [28]. (**B**) Multidimensional scaling plots gauging within- and across-order similarity of protein orthologous groups. Inset show color scheme for holometabolous orders. (**C**) Upset plot illustrating *C*. *felis* protein orthologous groups that intersect with other holometabolous insects. Inclusion criteria: one protein from at least one genome/order must be present. Yellow bar, 577 proteins found in all analyzed genomes. Inset, redrawn phylogeny estimation of Holometabola [2]; numbers indicate *C*. *felis* unique protein groups or higher-generic monophyletic groups (see **Additional file 5: Table S2**).

### Unique cat flea genome features

*C*. *felis* protein-encoding genes that failed to cluster with other Holometabola (4,282 sequences in 2,055 ortholog groups, **Fig. 3C**) potentially define flea-specific attributes. Elimination of divergent “holometabolan-like” proteins, identified with BLASTP against the nr database of NCBI, left 2,084 “unique” *C*. *felis* proteins (**Fig. 4A, Additional file 6: Table S3**). These include proteins lacking counterparts in the NCBI nr database (n=766), and proteins with either limited similarity to Holometabola or greater similarity to non-holometabolan taxa (n=1,318). Proteins comprising the latter set were assigned an array of functional annotations (GO, KEGG, InterPro, EC) and stand to guide efforts for deciphering flea-specific innovations (**Fig. 4B**, **Additional file 6: Table S3**).

**Fig. 4.**
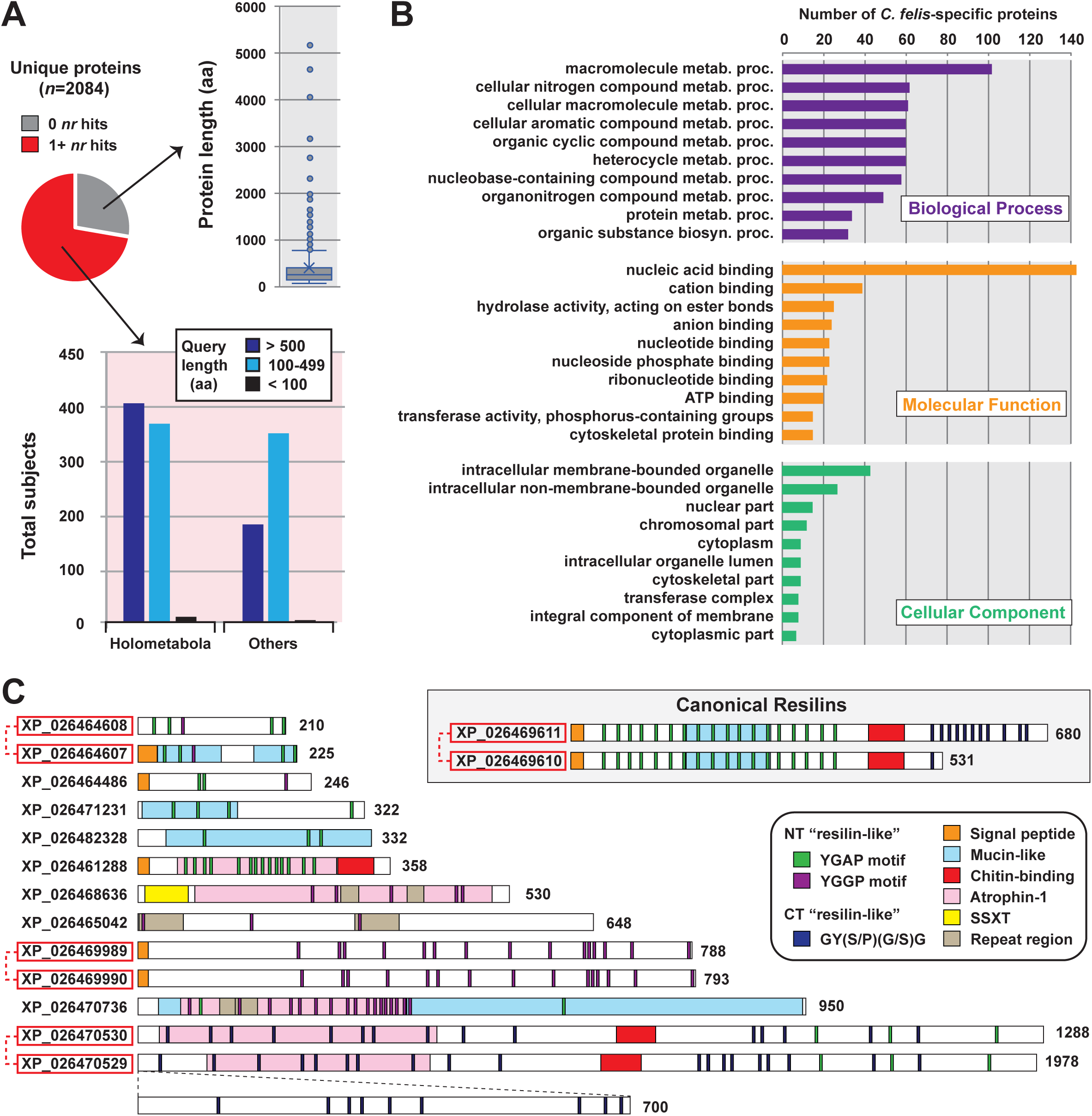
Identifying *C*. *felis*-specific genes. (**A**) *C*. *felis* proteins failing to cluster with counterparts in other holometabolan genomes were determined to lack (top) or possess limited similarity to (bottom) proteins from holometabolan or other genomes (bottom). (**B**) For 1,318 proteins, Gene Ontologies and Interpro domains were included in annotation and clustering into broad cellular function categories. (**C**) *C*. *felis* carries tandemly-arrayed resilin homologs (gray inset) as well as a cohort of other proteins containing resilin-like features. Red boxes indicate other tandemly-arrayed genes.

Two isoforms (A and B) of resilin, an elastomeric protein that provides soft rubber-elasticity to mechanically active organs and tissues, were previously identified in *C*. *felis* and proposed to underpin tarsal-mediated jumping [34]. Resilins typically have 1) highly repetitive Pro/Gly motifs that provide high flexibility, 2) key Tyr residues that facilitate intermolecular bonds between resilin polypeptides, and 3) a chitin-binding domain (CBD), though *C*. *felis* isoform B lacks the CBD [34, 35]. The *C*. *felis* assembly has two adjacent genes encoding resilins (gray box, **Fig. 4C**): the larger (680 aa) protein is more similar to both resilin A and B isoforms identified previously (>99 %ID), while the smaller (531 aa) protein is more divergent (98.8 %ID). These divergent resilins accentuate the observed CNV in *C*. *felis* and indicate additional genetic complexity behind flea jumping. Furthermore, a cohort of diverse proteins containing multiple resilin-like features and domains were identified, opening the door for future studies aiming to characterize the molecular mechanisms underpinning the great jumping ability of fleas.

### The *C*. *felis* microbiome: evidence for symbiosis and parasitism

Analysis of microbial-like Illumina reads revealed a bacterial dominance, primarily represented by *Proteobacteria* (**Fig. 5A**, **Additional file 7: Table S4**). Aside from the *Wolbachia* reads (discussed below), none of the bacterial taxa match to species previously detected in environmental [36, 37] or colony fleas [38]. Thus, a variable bacterial microbiome exists across geographically diverse fleas and is likely influenced by the presence of pathogens [38]. Strong matches to lepidopteran-associated *Chrysodeixis chalcites* nucleopolyhedrovirus and *Choristoneura occidentalis* granulovirus, as well as *Pandoravirus dulcis*, identify underappreciated viruses that may play important roles in the vectorial capacity of *C*. *felis*.

**Fig. 5.**
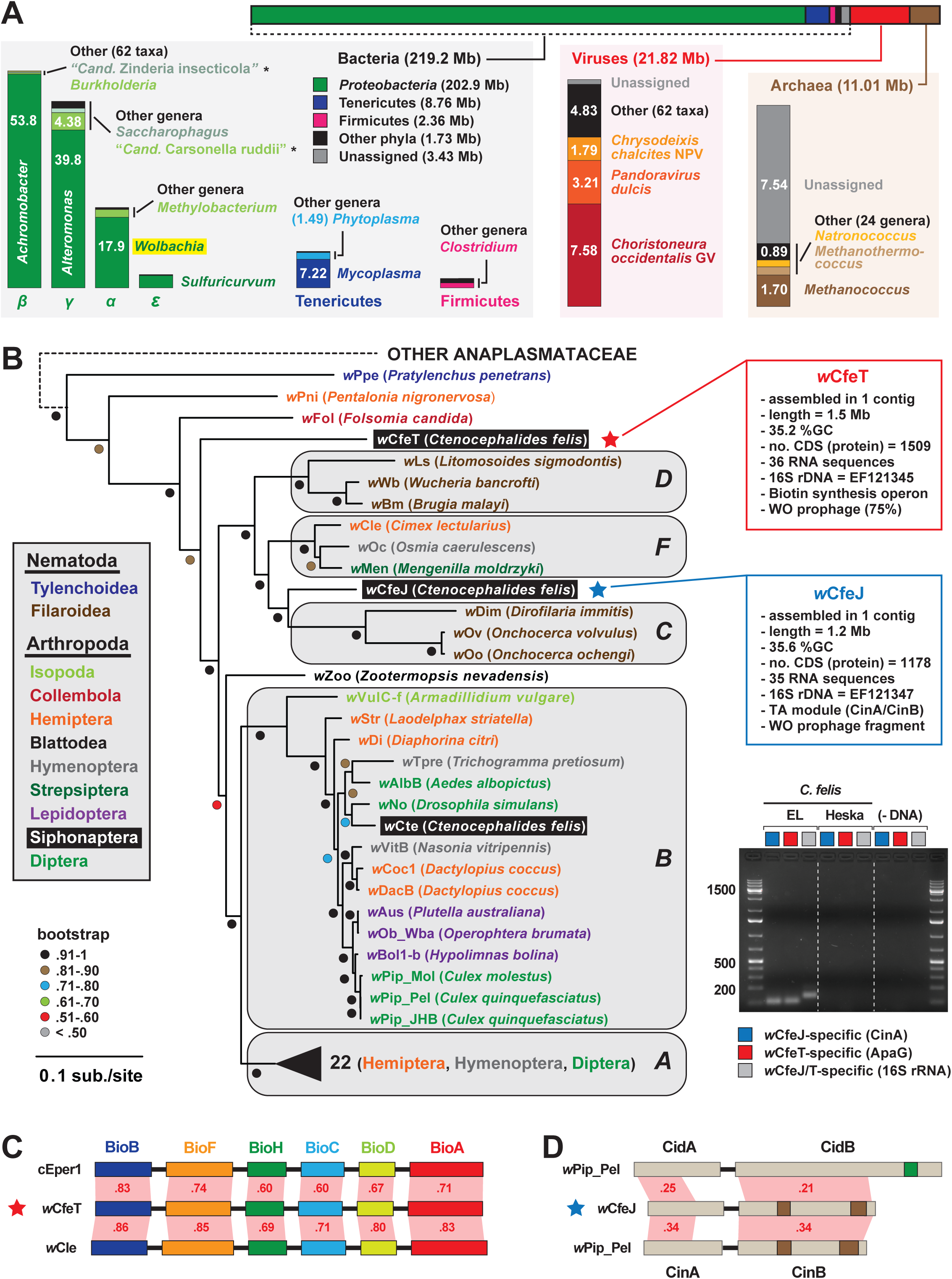
The microbiome of EL fleas. (**A**) Breakdown of the *C*. *felis* (EL fleas) microbiome. Bar at top graphically depicts the taxonomic distribution of non-flea Illumina reads across Bacteria, viruses and Archaea. Each group is further classified, with the major taxa (genus-level in most cases) and compiled read size (Mb) provided. Taxa with asterisks are AT-rich genomes that were later determined to match to *C*. *felis* mitochondrial reads. (**B**) *Wolbachia* genome-based phylogeny estimation. *Wolbachia* supergroups are within gray ellipses. *C*. *felis-*associated Wolbachiae are within black boxes. Red (*w*CfeT) and blue (*w*CfeJ) stars depict the two novel Wolbachiae infecting *C*. *felis*, with assembly information for each genome provided at right. Inset: color scheme for nematode and arthropod hosts. For tree estimation see **Methods**. Gel image (unaltered) depicts PCR results using 100ng of flea template DNA (quantified via nanodrop) in separate reactions with gene-specific primers. (**C**) *w*CfeT contains the unique biotin synthesis operon carried by certain obligately host-associated microbes. Schema follows our previous depiction of the unique *bio* gene order [44], with all proteins drawn to scale (as a reference, *w*CfeT BioB is 316 aa). Comparisons are made to the *bio* proteins of *Cardinium* endosymbiont of *Encarsia pergandiella* (cEper1, CCM10336-CCM10341) and *Wolbachia* endosymbiont of *Cimex lectularius* (*w*Cle, BAP00143-BAP00148). Red shading and numbers indicate % identity across pairwise protein alignments (blastp). (**D**) *w*CfeJ contains a CinA/B operon. Comparisons are made to the CidA/B (top, CAQ54390/1) and CinA/B (bottom) operons of *Wolbachia* endosymbiont of *Culex quinquefasciatus* Pel (wPip_Pel, CAQ54402/3). Green, CE clan protease; brown, PD-(D/E)XK nuclease. All proteins are drawn to scale (as a reference, *w*CfeJ CinB is 777 aa). Red shading and numbers indicate % identity across pairwise protein alignments (blastp).

Remarkably, two divergent *Wolbachia* genomes were assembled, circularized and annotated. Named *w*CfeT and *w*CfeJ, these novel strains were previously identified (using 16S rDNA) in a cat flea colony maintained at Louisiana State University [38–40], which historically has been replenished with EL fleas. Robust genome-based phylogeny estimation indicates *w*CfeT is similar to undescribed *C*. *felis*-associated strains that branch ancestrally to most other *Wolbachia* lineages [36, 41], while *w*CfeJ is similar to undescribed *C*. *felis*-associated strains closely related to *Wolbachia* supergroups C, D and F [42] (**Fig. 5B**; **Additional file 7: Table S4**). The substantial divergence of *w*CfeT and *w*CfeJ from a *Wolbachia* supergroup B strain infecting *C*. *felis* (*w*Cte) indicates a diversity of Wolbachiae capable of infecting cat fleas.

*w*CfeT and *w*CfeJ are notable for carrying segments of WO prophage, which are rarely present in genomes of Wolbachiae outside of supergroups A and B [43]. Further, each genome contains features that hint at contrasting relationships with *C*. *felis*. *w*CfeT carries the unique biotin synthesis operon (**Fig. 5C**), which was originally discovered in *Rickettsia buchneri* by us [44] and later identified in certain *Wolbachia* [45–47], *Cardinium* [48, 49] and *Legionella* [50] species. Given that some *Wolbachia* strains provide biotin to their insect hosts [45, 51], we posit that *w*CfeT has established an obligate mutualism with *C*. *felis* mediated by biotin-provisioning.

In contrast, *w*CfeJ appears to be a reproductive parasite, as it contains a toxin-antidote (TA) operon that is similar to the CinA/B TA operon of *w*Pip_Pel that induces cytoplasmic incompatibility (CI) in flies [52]. CinA/B operons are analogous to the CidA/B TA operons of *w*Mel and *w*Pip_Pel, which also induce CI in fly hosts [53–55], yet the CinB toxin harbors dual nuclease domains in place of the CidB deubiquitnase domain [56] (**Fig. 5D**). Given that the genomes of many *Wolbachia* reproductive parasites harbor diverse arrays of CinA/B-and CidA/B-like operons [56, 57], wCfeJ’s CinA/B TA operon might function in CI or some other form of reproductive parasitism. Quizzically, the co-occurrence of *w*CfeJ and *w*CfeT in individual fleas (gel image in **Fig. 5B**) indicates dual forces (mutualism, parasitism) that potentially drive their infection in EL fleas.

## Discussion

We set out to generate a genome sequence for the cat flea, a surprisingly absent resource for comparative arthropod genomics and vector biology. Our efforts to generate a *C*. *felis* assembly brought forth an unexpected finding, namely that no two cat fleas share the same genome sequence. We provide multiple lines of evidence supporting flea genomes in flux (**Table 1**).

**Table 1.**
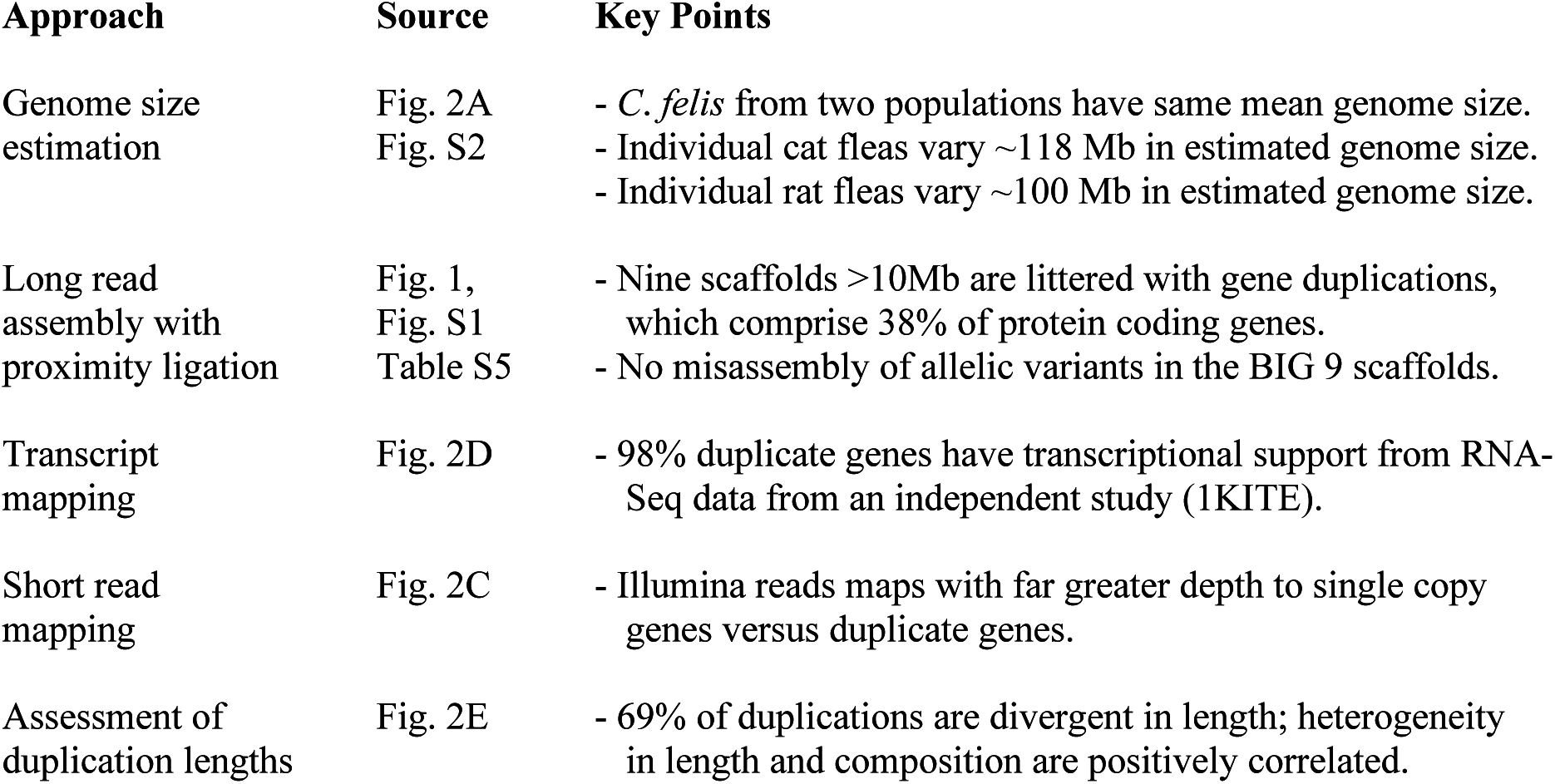
Evidence Supporting Extensive Gene Duplication in Cat Fleas.

First, genome size estimations for over two dozen individual cat fleas from the EL colony revealed over 150 Mb variation, a result consistent with prior genome size estimates for *C*. *felis* from a different colony as well as rat fleas. Second, our haplotig-resolved assembly identified rampant gene duplication throughout the genome. Third, RNA-Seq data from an independent colony confirmed the pervasive gene duplication. Finally, ∼70% of gene duplications are not comparable in length, indicating active gene expansion and contraction. Since transposons and other repeat elements are relatively sparse in *C. felis* and cannot account for such rampant CNV, and given that no individual flea genome size was estimated to be larger than our BIG9 assembly, we posit that unequal crossing over and gene conversion continually create and eliminate large linear stretches of DNA to keep the *C*. *felis* genome in a fluctuating continuum. We favor this hypothesis over an ancient whole genome duplication event in Siphonaptera provided that the majority of these duplications are tandem or proximal.

Ramifications of a genome in flux are readily identifiable. First, as gene duplication is a major source of genetic novelty, extensive CNV likely affords *C*. *felis* with a dynamic platform for innovation, allowing it to outpace gene-targeting pest control measures. Second, extensive CNV will complicate standard normalization procedures utilized in comparative transcriptomics analysis, requiring a more nuanced interpretation of standard metrics that are based on gene length (i.e. RPKM, TPM, etc.). Furthermore, achieving high confidence with read-mapping to cognate genes will be difficult in the face of neofunctionalization, subfunctionalization and early pseudogenization, as well as dosage-based regulation of duplicate genes. Third, genetic markers typically utilized for evolutionary analyses (e.g., phylotyping, population genetics, phylogeography [58]) may yield erroneous results when applied to *C*. *felis* and related *Ctenocephalides* species if targeted to regions of CNV (and particularly neofunctionalization). Finally, as a *C*. *felis* chromosome-level genome assembly was only attainable by coupling Illumina and PacBio sequencing with Hi-C scaffolding techniques, short-read based sequencing strategies will be inadequate for other organisms with high CNV. The ability of the BIG9 assembly to serve as a reference genome in future short-read based sequencing efforts for other cat fleas will be determined. Moving forward, newly developed low-input protocols for PacBio sequencing will allow us to query individual fleas to robustly assess the degree of gene duplication.

Excessive CNV in *C*. *felis*, and likely all Siphonaptera, requires the determination of the genetic mechanisms at play. Why extreme gene duplication, when predicted across arthropods using genomic and transcriptomic data [59], was not previously detected in fleas is unclear. Excessive CNV aside, our study provides the first genome sequence for Siphonaptera, which will substantially inform comparative studies on insect vectors of human disease. Furthermore, newly-identified symbiotic (*w*CfeT) and parasitic (*w*CfeJ) *Wolbachia* will be paramount to efforts for biocontrol of pathogens transmitted by cat fleas. The accrued resources and knowledge from our study are timely. A drastic rise of murine typhus cases alone in Southern California [60] and Galveston, Texas [61], which are directly attributable to fleas associated with increasing population sizes of rodents and opossums, requires immediate and re-focused efforts to combat this serious and underappreciated risk to human health.

## Conclusion

Fleas are parasitic insects that can transmit many serious pathogens (i.e. bubonic plague, endemic and murine typhus). The lack of flea genome assemblies has hindered research, especially comparisons to other disease vectors. Here we combined Illumina and PacBio sequencing with Hi-C scaffolding techniques to generate a chromosome-level genome assembly for the cat flea, *Ctenocephalides felis*. Our work has revealed a genome characterized by inordinate copy number variation (∼38% of proteins) and a broad range of genome size estimates (433-551 Mb) for individual fleas, suggesting a bizarre genome in flux. Surprisingly, the flea genome exhibits neither inflation due to rampant gene duplication nor reduction due to their parasitic lifestyle. Based on these results, as well as the nature and distribution of the gene duplications themselves, we posit a dual mechanism of unequal crossing-over and gene conversion may underpin this genome variability, although the biological significance remains to be explored. Coupled with paradoxical co-infection with novel *Wolbachia* endosymbionts and reproductive parasites, these oddities highlight a unique and underappreciated human disease vector.

## Methods

### Experimental design

This study was undertaken to generate a high-quality reference genome assembly and annotation for the cat flea, *C. felis*, and represents the first sequenced genome for a member of Order Siphonaptera. Our approach leveraged a combination of long-read PacBio sequencing, short-read Illumina sequencing, and Hi-C (Chicago and HiRise) data to construct a chromosome-level assembly; RNA-seq data and BLAST2GO classifications to assist in gene model prediction and annotation; sequence mapping to address assembly fragmentation and short scaffolds (<1Mb); and ortholog group construction to explore a genetic basis for the cat flea’s parasitic lifestyle. Gene duplications were confirmed via orthogonal approaches, including genome size estimates of individual fleas, gene-based read coverage calculations, genomic distance between duplications, and correlation between duplications and repeat elements or contig boundaries.

### Genome Sequencing and Assembly

Newly emerged (August 2017), unfed female *C. felis* (n = 250) from Elward Laboratories (EL; Soquel, CA) were surface-sterilized for 5 min in 10% NaClO followed by 5 min in 70% C_2_H_5_OH and 3X washes with sterile phosphate-buffered saline. Fleas were flash-frozen in liquid N_2_ and ground to powder with sterile mortar and pestle. High-molecular weight DNA was extracted using the MagAttract HMW DNA Kit (Qiagen; Venlo, Netherlands), quantified using a Qubit 3.0 fluorimeter (Thermo-Fisher Scientific; Waltham, MA), and assessed for quality on a 1.5% agarose gel. DNA (50 µg) was submitted to the Institute for Genome Sciences (University of Maryland) for size-selection and preparation of sequencing libraries. Libraries were sequenced on 12 SMRT cells of a PacBio Sequel (Pacific Biosciences; Menlo Park, CA), generating 7,239,750 reads (46.7 Gb total). Raw reads were corrected, trimmed, and assembled into 16,622 contigs with Canu v1.5 in “pacbio-raw” mode, using an estimated genome size of 465 Mb [30]. A second group of newly emerged (January 2016), unfed female EL fleas (n=100) was surface-sterilized and homogenized as above, and genomic DNA extracted using the QIAgen DNeasy® Blood and Tissue Kit (QIAgen, Hilden, Germany). DNA was submitted to the WVU Genomics Core for the preparation of a paired-end 250bp sequencing library with an average insert size of 500bp. The library was sequenced on 4 lanes of an Illumina HiSeq 1500 (Illumina Inc.; San Diego, CA), generating 450,132,548 reads which were subsequently trimmed to remove adapters and filtered for length and quality using FASTX-Toolkit v0.0.14 (available from http://hannonlab.cshl.edu/fastx_toolkit/). These short read data were used to polish the Canu assembly with Pilon v1.1.6 in “fix-all” mode [62], and to determine the composition of the *C. felis* microbiome (see below). Haplotigs in the polished contigs were resolved using purge_haplotigs [31] with coverage settings of 5 (low), 65 (mid), and 180 (high). A third group of newly-emerged (Feburary 2018), unfed female EL fleas (n = 200) were surface-sterilized as above, frozen at −80°C, and submitted for Chicago and Dovetail Hi-C proximity ligation (Dovetail Genomics, Santa Cruz, CA) [63] using the polished Canu assembly as a reference. The resulting scaffolded assembly (3,926 scaffolds) was subjected to removal of microbial sequences as described in the next section.

### Genome Decontamination

A comparative BLAST-based pipeline slightly modified from our prior work [64] was used to identify and remove microbial scaffolds before annotation. Briefly, polished contigs were queried using BLASTP v2.2.31 against two custom databases derived from the nr database at NCBI (accessed July 2018): (1) all eukaryotic sequences (eukDB), and (2) combined archaeal, bacterial, and viral sequences (abvDB). For each query, the top five unique subject matches (by bitscore) in each database were pooled and scored according to a comparative sequence similarity measure, S_m_:

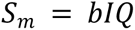

where *b* is the bitscore of the match; *I* is the percent identity; and *Q* is the percent aligned based on the longer of the two sequences. The top 5 scoring matches from the pooled lists of subjects were used to calculate a comparative rank score *C* for each individual query *q* against each database *d*:

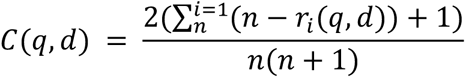

where *r_i_(q,d)* is the rank of subject *i* for query *q* against database *d*. For example, if all of the top *n* matches for query *q* are in eukDB then *C(q,*eukDB) = 1; conversely, if none of the top *n* matches are in database abvDB then *C(q,*abvDB) = 0. Finally, each query *q* was scored according to a comparative pairwise score *P* between 1 purely eukaryotic) and −1 (purely microbial):

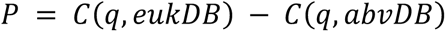

Scaffolds that contained no contigs with *P* > 0.3 (n = 183), including 5 *Wolbachia*-like scaffolds, were classified “not eukaryotic” and set aside. Scaffolds that contained contigs with a range of P scores (n = 32) were manually inspected to identify and remove scaffolds arising from misassembly or contamination (n = 10). The remaining scaffolds (n = 3,733) comprised the initial draft assembly for *C. felis* and were deposited in NCBI under the accession ID GCF_003426905.1.

### Genome Annotation

Assembled and decontaminated scaffolds were annotated with NCBI Eukaryotic Genome Annotation Pipeline (EGAP) v8.1 (https://www.ncbi.nlm.nih.gov/books/NBK143764/). To facilitate gene model prediction, we generated RNA-seq data from 6 biological replicates of pooled *C. felis* females (Heska Corporation, Fort Collins, CO). Briefly, total RNA was isolated and submitted to the WVU Genomics Core for the preparation of paired-end, 100 bp sequencing libraries using ScriptSeq Complete Gold Kit for Epidemiology (Illumina Inc., San Diego, CA). Barcoded libraries were sequenced on 2 lanes of an Illumina HiSeq 1500 in High Throughput mode, yielding approximately 26 million reads per sample (Q > 30). Raw sequencing reads from all 6 samples were deposited in NCBI under the BioProject accession PRJNA484943. In addition to these data, the EGAP pipeline also integrated previously-published *C. felis* expression data from the 1KITE project (accession SRX314844; [2]) and an unrelated EST library (Biosample accession SAMN00161855). The final set of annotations is available as “Ctenocephalides felis Annotation Release 100” at the NCBI.

### Genome Completeness and Deflation

The distribution of scaffold lengths in our assembly, together with the relatively large number of fleas in our sequenced pool, warranted evaluating short scaffolds as possible sources of genomic heterogeneity among individual fleas. To address this possibility, assembly scaffolds shorter than 1 Mb (n = 3,724) were mapped to scaffolds larger than 1 Mb (n = 9; the BIG9) with BWA-MEM v0.7.12 [65] using default parameters (**Additional file 1: Fig. S1A**). Additionally, genome completeness of the full assembly compared to just the BIG9 scaffolds was assessed with Benchmarking Using Single Copy Orthologs (BUSCO) v3.0.2 [28] in “protein” mode, using the *eukaryota_odb9*, *arthropoda_odb9*, and *insecta_odb9* data sets (**Additional file 1: Fig. S1B**). Isoforms were removed before BUSCO analysis by identifying CDSs that derived from the same protein-coding gene and removing all but the longest sequence.

### Assessing the Extent of Gene Duplication

Proteins encoded on the BIG9 scaffolds (n = 16,518) were queried against themselves with BLASTP v2.2.31 using default parameters. Pairs of unique sequences that met or exceeded a given amino acid percent identity (%ID) threshold over at least 80% of the query length were binned together. Bins of sequence pairs that shared at least one sequence in common were subsequently merged into clusters. Isoforms were removed after clustering by identifying CDSs in a cluster that derived from the same protein-coding gene and removing all but the longest sequence. This process was used to generate cluster sets at integer %ID thresholds from 90% to 100%. These duplicate protein-encoding genes were then mapped onto each of the BIG9 scaffolds using Circos [66] (**Additional file 1: Fig. S1C-K**). Cluster diameters were calculated as the number of non-cluster genes that lie between the edges of the cluster (*i.e.*, the two cluster genes that are farthest apart on the scaffold) (**Additional file 1: Fig. S1L**). Clusters that span multiple scaffolds (mapped across all BIG9 scaffolds in **Additional file 1: Fig. S1M**) defy an accurate calculation of diameter and were assigned a cluster diameter of −1. In order to estimate the fraction of our assembly comprising gene duplications, cluster coverages (by %ID threshold) were calculated in three ways. First, the *coverage by CDS* was estimated by comparing the number of single-copy (protein-encoding) genes to the total number of clusters; the latter number is assumed to represent a theoretical set of minimal “seed” sequences. Second, the *coverage by gene length* was calculated as the total number of nucleotides encoding the proteins in each cluster (including introns and exons) minus the mean gene length (to account for a hypothetical “ancestor” gene). Finally, the *coverage by genome region* was estimated by adding i*(n-1) to each calculation of coverage by gene length, where n is the number of genes in the cluster and i is the mean intergenic length across all BIG9 scaffolds (17,344 nt). In order to assess possible enrichment of cellular functions among duplicated genes, clusters at the 90% ID level were compared to the remaining BIG9 proteins by Fisher’s Exact Test (corrected for multiple testing) which is integrated into the FatiGO package of BLAST2GO (see section “*Functional Classification of* C. felis *Proteins*” below). GO categories were reduced to their most specific terms whenever possible.

### Length Variation Within Gene Duplication Clusters

Variability in intra-cluster CDS length was assessed in two ways. First, the length of each CDS in a cluster was compared to the longest CDS of the cluster, and the proportion of clusters with any truncation (>1 AA) was calculated for each integer %ID threshold between 90 and 100% ID. Second, the mean and distribution of length differences (i.e., the extent of truncation) was calculated across all clusters for each integer %ID threshold between 90 and 100% ID.

### Analysis of Repeat Regions

The extent and composition of repeat elements in the *C. felis* genome were assessed in two ways. First, proteins annotated in the GO category “DNA Integration GO:0015074” (including retrotransposons) were extracted, plotted by genomic coordinate on each BIG9 scaffold, and assessed for co-localization either with gene duplicates (see above) or near the ends of scaffolds (**Additional file 1: Fig. S1N**). Second, repeat elements were identified on the BIG9 scaffolds with RepeatMasker v4.0.9 (available from http://www.repeatmasker.org/) in “RMBlast” mode (species “holometabola”), using Tandem Repeat Finder v4.0.9 and the Repbase RepeatMasker (October 2018) and Dfam 3.0 databases (**Additional file 1: Fig. S1O**).

### Codon Usage and tRNA Gene Family Analysis

Given the relatively large number of tRNA genes in our assembly, and the AT richness of our genome, we were interested in exploring connections between tRNA gene frequencies and codon usage. To this end, tRNA gene abundance on BIG9 scaffolds (n = 4,358) was determined by binning genes into families according to their cognate amino acid and calculating the percent of each family compared to the total number of tRNA genes (**Additional file 1: Fig. S1P**). A similar approach was taken to quantify tRNA gene abundance by anticodon. TA richness of each anticodon was subsequently calculated as the percent of A+T bases in the anticodon corrected for the size of the tRNA family. Codon usage was calculated as the percent of total codons using the coding sequences for genes on the BIG9 scaffolds, with isoforms removed as described previously (**Additional file 1: Fig. S1Q**).

### Functional Classification of *C. felis* Proteins

Protein sequences encoded on the BIG9 scaffolds (n = 16,518) were queried with BLASTP v2.2.31 against the nr database of NCBI (accessed July 2018) using a maximum e-value threshold of 0.1. The top 20 matches to each *C. felis* sequence were used to annotate queries with Gene Ontology (GO) categories, Enzyme Classification (EC) codes, and protein domain information using BLAST2GO v1.4.4 [67] under default parameters. A local instance of the GO database (updated February 2019) was used for GO classification, and the online version of InterPro (accessed April 2019) was used for domain discovery, including InterPro, PFAM, SMART, PANTHER, PHOBIUS, and GENE3D domains; PROSITE profiles; SignalP-TM (signal peptide) domains; and TMHMM (transmembrane helix) domains. InterPro data was used to refine GO annotations whenever possible (**Additional file 2: Table S1**). A subset of *C. felis* proteins (n = 153) classified as “DNA repair” (GO:0006281) was identified and all child GO terms of these proteins tabulated (**Additional file 2: Table S1**). Assuming a linear relationship between genome size and number of repair genes [68], we estimate *C. felis* has an enriched repertoire closer to that of a 3 Gb genome.

### Genome Size Estimation

Estimations for flea genome size largely followed previously reported approaches [69]. For *C*. *felis* individuals, 1/20 of the flea head was combined with two standards: 1/20 of the head of a female (YW) *Drosophila melanogaster* (1C = 175 Mbp) and 1/20 of the head of a lab strain *D*. *virilis* female (1C = 328). The tissues were placed in 1ml of cold Galbraith buffer and ground to release nuclei in a 2ml Kontes Dounce, using 15 strokes of the “A” pestle at a rate of three strokes every two seconds. The resulting solution was strained through a 45µ filter, stained for 3 hours in the dark at 4°C with 25µl of propidium iodide, then scored for total red fluorescence using a Beckman-Coulter CytoFLEX flow cytometer. The average channel number of the 2C nuclei of the sample and standards were determined using the CytExpert statistical software. Briefly, the amount of DNA was estimated as the ratio of the average red fluorescence of the sample to the average red fluorescence of the standard multiplied by the amount of DNA (in Mbp) of the standard. The estimates from the two standards were averaged. At least 500 nuclei were counted in each sample and standard peak. The coefficients of variation (CV) for all peaks were < 2.0. Fluorescence activation and gating based on scatter were used to include in each peak only intact red fluorescent nuclei free of associated cytoplasmic or broken nuclear tags. Histograms generated for the largest and smallest determined genome sizes show the minimal change in position for the two standards, demonstrating the significant change in the relative fluorescence (average 2C channel number) between *C*. *felis* individuals (**Additional file 3: Fig. S2**).

### Characterizing Copy Number Variation

In order to test the hypothesis that our genome assembly represents an agglomeration of individuals with different levels of gene duplication, we used minimap2 [70] to map our short-read sequence data against the full scaffolded assembly. After extracting the mapped reads with samtools v0.1.19 [71], including primary and alternative mapping loci, a vector of sequence depth (in bases) per position was generated with the genomecov function of bedtools v2.25.0 [72]. Mean depths for all 16,518 protein-coding genes on the BIG9 scaffolds were calculated as total bases covering each gene divided by gene length. Finally, the mean depth across all duplicated genes was compared to the mean depth across all single-copy genes using a Student’s t-test.

To evaluate the extent of gene duplication across different *C. felis* populations, reads from the 1KITE transcriptome sequencing project (NCBI Sequence Read Archive accession SRR921588) were mapped to the 3,733 scaffolds from our assembly using HISAT2 v2.1.0 [73] under the --dta and --no_unal options. Mapped reads were sorted with samtools and abundance per gene calculated as transcripts per million reads (TPM) using stringtie v1.3.4d [73]. TPM values were binned and plotted against the number of duplicated (90% aa ID or higher) and single-copy genes in the BIG9 assembly.

### Comparative Genomics

Protein sequences (n=1,077,182) for 51 sequenced holometabolan genomes were downloaded directly from NCBI (n=47) or VectorBase (n=3) or sequenced here (n=1). Isoforms were removed before analysis wherever possible, by identifying CDSs that derived from the same protein-coding gene and removing all but the longest CDS. Genome completeness was estimated with BUSCO v3.0.2 in “protein” mode, using the *insecta_odb9* data set. Ortholog groups (OGs; n=50,118) were constructed in three sequential phases: 1) CD-HIT v4.7 [74] in accurate mode (-g 1) was used to cluster sequences at 50% ID; 2) PSI-CD-HIT (accurate mode, local identity, alignment coverage minimum of 0.8) was used to cluster sequences at 25% ID; 3) clusters were merged using clstr_rev.pl (part of the CD-HIT package). Proteins from *C. felis* that did not cluster into any OG (n=4,282) were queried with BLASTP v2.2.31 against the nr database of NCBI (accessed July 2018). Queries (n=2,170) with a top hit to any Holometabola taxon, at a minimum %ID of 25% and query alignment of 80%, were manually added to the original set of ortholog groups where possible (n=2,142) or set aside where not (n=28). The remaining queries with at least one match in nr (n=1,318) were grouped by GO category level 4 and manually inspected; these included queries with top hits to Holometabolan taxa that did not meet the minimum %ID or query coverage thresholds. Finally, *C. felis* proteins with no match in nr (n=766) were binned by query length. These last two sets (n=2,084) comprise the set of proteins unique to *C. felis* among all other Holometabola (**Additional file 6: Table S3**). Congruence between OG clusters and taxonomy was determined by calculating a distance (Euclidean) between each pair of taxa based on the number of shared OGs. The resulting matrix was scaled by classic multidimensional scaling with the cmdscale function of R v3.5.1 [75], and visualized using the ggplot package in R. Finally, pan-genomes were calculated for several key subsets of Holometabola: 1) *C. felis* alone (Siphonaptera); 2) Antliophora (Siphonaptera and Diptera); 3) Panorpida (Siphoanptera, Diptera, and Coleoptera); 4) all taxa except Hymenoptera; and 5) all Holometabola (**Additional file 5: Table S2**). In order to account for differences in genome assembly quality and taxon sampling bias, we define the pan-genome here as the set of all OGs that contain at least one protein *from at least one taxon* in a given order. These intersections were visualized as upset plots using UpSetR v1.3.3 [76]. Intersections of various holometabolous taxa that lack *C*. *felis* were computed to gain insight on possible reductive evolution in fleas (**Additional file 4: Fig. S3**, **Additional file 5: Table S2**).

### Microbiome Composition

A composite *C. felis* microbiome was estimated using Kraken Metagenomics-X v1.0.0 [77], part of the Illumina BaseSpace toolkit. Briefly, 105,256,391 PE250 reads from our short read data set were mapped against the Mini-Kraken reference set (12-08-2014 version), resulting in 2,390,314 microbial reads (2.27%) that were subsequently assigned to best possible taxonomy (**Additional file 7: Table S4**).

### Assembly of *Wolbachia* Endosymbiont Genomes

Corrected reads from the Canu assembly of *C. felis* were recruited using BWA-MEM v0.7.12 (default settings) to a set of concatenated closed *Wolbachia* genome sequences (n=15) downloaded from NCBI (accessed February 2018). Reads that mapped successfully were extracted with samtools v0.1.19 and assembled separately into seed contigs (n=22) with Canu v1.5 using default settings. Gene models on these seed contigs were predicted using the Rapid Annotation of Subsystems Technology (RAST) v2.0 server [78], yielding two small subunit (16S) ribosomal genes that were queried with BLASTN against the nr database of NCBI to confirm the presence of two distinct *Wolbachia* strains. Seed contigs were further analyzed by %GC and top BLASTN matches in the nr database of NCBI, and binned into three groups: *C. felis* mitochondrial (n=1), *C. felis* genomic (n=6), and *Wolbachia*-like (n=15) contigs. The *Wolbachia*-like contigs were subsequently queried with BLASTN against the full *C. felis* assembly (before decontamination). A single *Wolbachia*-like contig (tig00000005; wCfeJ) containing one of the two distinct 16S genes was retrieved intact from the full assembly. It was removed from the primary assembly and manually closed by aligning the contig ends with BLASTN. Gaps in the aligned regions were resolved by mapping our short read data to the contig with BWA-MEM (default settings) and manually inspecting the read pileups. Six additional contigs were also retrieved intact from the full assembly; these were likewise removed and manually stitched together using end-alignment and short read polishing, resulting in a second closed *Wolbachia* genome (*w*CfeT). The remaining *Wolbachia*-like contigs (n=8) were found to be fractions of much longer flea-like contigs; these were left in the primary *C. felis* assembly. Both *w*CfeJ and *w*CfeT sequences were submitted to the RAST v2.0 server for gene model prediction and functional annotation.

### Phylogenomics of *Wolbachia* Endosymbionts

Protein sequences (n=66,811) for 53 sequenced *Wolbachia* genomes plus 5 additional Anaplasmataceae (*Neorickettsia helminthoeca* str. Oregon, *Anaplasma centrale* Israel, *A. marginale* Florida, *Ehrlichia chaffeensis* Arkansas, and E. *ruminantium Gardel*) were either downloaded directly from NCBI (n=30), retrieved as genome sequences from the NCBI Assembly database (n=13), contributed via personal communication (n=8; Michael Gerth, Oxford Brookes University), or sequenced here (n=2) (**Additional file 7: Table S4**). For genomes lacking functional annotations (n=15), gene models were predicted using the RAST v2.0 server (n=12) or GeneMarkS-2 v1.10_1.07 (n=3; [79]). Ortholog groups (n=2,750) were subsequently constructed using FastOrtho, an in-house version of OrthoMCL [80], using an expect threshold of 0.01, percent identity threshold of 30%, and percent match length threshold of 50% for ortholog inclusion. A subset of single-copy families (n=47) conserved across at least 52 of the 58 genomes were independently aligned with MUSCLE v3.8.31 [81] using default parameters, and regions of poor alignment were masked with trimal v1.4.rev15 [82] using the “automated1” option. All modified alignments were concatenated into a single data set (10,027 positions) for phylogeny estimation using RAxML v8.2.4 [83], under the gamma model of rate heterogeneity and estimation of the proportion of invariant sites. Branch support was assessed with 1,000 pseudo-replications. Final ML optimization likelihood was -183020.639712.

### Confirmation of the presence of *Wolbachia* in *C. felis*

To assess the distribution of *w*CfeJ and *w*CfeT in *C. felis*, individual fleas from the sequenced strain (EL) and a separate colony (Heska) not known to be infected with *Wolbachia* were pooled (n=5) by sex and colony, surface-sterilized with 70% ethanol, flash-frozen, and ground in liquid N_2_. Genomic DNA was extracted using the GeneJET Genomic DNA Extraction Kit (Thermo-Fisher Scientific; Waltham, MA), eluted twice in 50µl of PCR-grade H_2_O, and quantified by spectrophotometry with a Nanodrop 2000 (Thermo-Fisher Scientific; Waltham, MA). 100ng of DNA from each pool was used as template in separate 25 µl PCR reactions using AmpliTaq Gold 360 (Thermo-Fisher Scientific; Waltham, MA) and primer pairs (400 nmoles each) specific for: 1) a 76nt fragment of the *cinA* gene specific to *w*CfeJ (Fwd: 5’-AGCAACACCAACATGCGATT-3’; Rev: 5’-GAACCCCAGAGTTGGAAGGG-3’); 2) a 75nt fragment of the *apaG* gene specific to *w*CfeT (Fwd: 5’-GCCGTCACTGGCAGGTAATA-3’; Rev: 5’-GCTGTTCTCCAATAACGCCA-3’); or 3) a 122nt fragment of *Wolbachia* 16S rDNA (Fwd: 5’-CGGTGAATACGTTCTCGGGTY-3’; Rev: 5’-CACCCCAGTCACTGATCCC-3’). Primer specificities were confirmed with BLASTN against both the *C. felis* assembly and the nr database of NCBI (accessed June 2018). Reaction conditions were identical for all primer sets: initial denaturation at 95°C for 10 min, followed by 40 cycles of 95°C for 30 sec, 60°C for 30 sec, and 72°C for 30 sec, and a final extension at 72°C for 7 min. Products were run on a 2% agarose gel and visualized with SmartGlow Pre Stain (Accuris Instruments; Edison, NJ). Primers were tested before use by quantitative real-time PCR on a CFX Connect (Bio-Rad Laboratories; Hercules, CA).

### Statistical Analysis

Statistical analyses were carried out in R v3.5.1. Mean coverages across duplicated (n=7852) and single-copy (n=7061) genes at the 90% ID threshold were compared for significance using a Welch Two Sample t-test (unpaired, two-tailed) with 12,930 degrees of freedom and a p-value < 2.2×10^−16^. Mean coverage of duplicated genes at %ID thresholds from 85-100% were compared for significance using one-way Analysis of Variance (ANOVA) with 15 degrees of freedom and a p-value = 0.2. A similar ANOVA was used to compare single-copy genes at 85-100% ID thresholds, with a p-value < 2.2×10^−16^.

### Data and Scripts

Data generated for this project that is not published elsewhere, including BLAST2GO annotations and OG assignments, as well as custom analysis scripts, are provided on GitHub in the “cfelis_genome” repository available at https://www.github.com/wvuvectors/cfelis_genome.

## Declarations

### Ethics approval and consent to participate

Not applicable.

### Consent for publication

Not applicable.

### Availability of data and materials

All of the sequence data generated for this work are available at the NCBI under Bioproject accessions PRJNA489463 (genome sequence and annotation) and PRJNA484941 (RNA-seq data used to support annotation). Additional tables with GO annotations, ortholog groups, and microbiome data, as well as scripts used to generate data visualizations can be accessed at https://www.github.com/wvuvectors/cfelis_genome. Sequences for *w*CfeT and *w*CfeJ are available on NCBI under Bioproject PRJNA622233.

### Authors’ contributions

TPD, VIV, JJG, KRM, and AFA conceived this study and developed the overall experimental framework. TPD, VIV, and KRM isolated genomic DNA from fleas. MLG, KER-B, and MSR performed flea dissections and isolated RNA from flea midgut tissues. TPD, VIV, JJG, DH, and CGE conceptualized the strategies for assembly and annotation. TPD, VIV, and JJG performed quality control at various stages of the assembly, executed the overall analyses of flea annotation, performed the phylogenomics analyses, and analyzed the flea microbiome. JSJ estimated genome sizes for individual cat and rat fleas. All authors contributed to writing and review of the final manuscript, with TPD, VIV, JJG, JSJ, KRM, and AFA playing the key roles. All authors read and approved the final manuscript.

## Acknowledgements

We would like to thank Luke Tallon and Lisa Sadzewicz (Institute for Genome Sciences, University of Maryland, Baltimore) for assistance with PacBio sequencing, Ryan Percifield (Genomics Core, West Virginia University) for Illumina sequencing, and Mark Daly, Elizabeth Fournier, Shaune Hall, and Thomas Swale (Dovetail Genomics) for facilitating Hi-C assembly. We acknowledge the generous gift of unpublished *Wolbachia* endosymbiont of *Zootermopsis nevadensis* (wZoo), *Wolbachia* endosymbiont of *Mengenilla moldrzyki* (wMen), and *Wolbachia* endosymbiont of *Ctenocephalides felis* (wCte) contigs from Michael Gerth (University of Liverpool). We thank Dr. Joe Hinnebusch (National Institute of Allergy and Infectious Diseases) for providing the rat fleas used for genome size estimation. We are grateful to Dr. John Beckmann (Auburn University) for critical discussion regarding *Wolbachia* biology and Magda Beier-Sexton (University of Maryland, Baltimore) for administrative support.

---

**Additional file 1: Fig. S1.** Assessing assembly fragmentation, gene duplication and repeat elements within the *C*. *felis* assembly. (**A**) Evaluating assembly fragmentation via mapping of scaffolds shorter than 1 Mb (n = 3,724) to scaffolds larger than 1 Mb (n = 9, “BIG9 scaffolds”). All but 2 short scaffolds mapped to a BIG9 scaffold at least once; confidence intervals are based on the probability of mapping to a single unique location. (**B**). Assessing the “genome completeness” of the *C*. *felis* full assembly and BIG9 scaffolds through comparison to eukaryote, arthropod and insect BUSCOs. (**C**) Tandem and proximal duplicate gene locations on BIG9 scaffold 1, (**D**) BIG9 scaffold 2, (**E**) BIG9 scaffold 3, (**F**) BIG9 scaffold 4, (**G**) BIG9 scaffold 5, (**H**) BIG9 scaffold 6, (**I**) BIG9 scaffold 7, (**J**) BIG9 scaffold 8, (**K**) BIG9 scaffold 9. (**L**) Duplications by proximity. Only true duplications (n=2) are shown. Red bars (*) depict “dispersed” clusters that span multiple scaffolds. (**M**) Dispersed duplicate gene locations across BIG9 scaffolds. (**N**) Distribution across BIG9 scaffolds of *C*. *felis* proteins annotated as “DNA integration” (GO:0015074, see **Additional file 2: Table S1** for specific accession numbers) and their relation to gene duplications. (**O**) Compilation of retroelements, DNA transposons and other repeat elements predicted across the BIG9 scaffolds. Overall totals are highlighted yellow. (**P**) tRNA gene abundances and (**Q**) codon usage/amino acid for select Holometabola.

**Additional file 2: Table S1.** Functional predictions and enrichment analysis of *C*. *felis* proteins.

<click for link to Table S1>

**Additional file 3: Fig. S2.** Representative histograms produced by flow cytometry showing the peak positions of the 2C nuclei of *Drosophila melanogaster* (left) and *D. virilis* (center) female standards, and individual *C. felis* females (right) from the sequenced EL strain. (**A**) A 434 Mb flea. (**B**) A 553 Mb flea. All peaks have CV < 1.5 and > 500 nuclei under the statistical gates (red lines spanning the 2C peaks).

**Additional file 4: Fig. S3.** Phylogenomics analysis of select Holometabola. (**A**) Assessment of holometabolan accessory genomes. (**B**) *Top:* Identification of conserved protein families present in select taxa from each holometabolan order but absent from *C*. *felis*. *Bottom:* Protein families conserved across all sequenced holometabolan genomes except *C. felis* (see **Additional file 5: Table S2**). Four assemblies were identified as particularly patchy (*Oryctes borbonicus*, *Operophtera brumata*, *Heliothis virescens*, and *Plutella xylostella*) and 100% conservation (”perfect”) was also relaxed to exclude these taxa. Inset, redrawn phylogeny estimation of Holometabola [2].

**Additional file 5: Table S2.** Pan-genomes across sequenced Holometabola.

<click for link to Table S2>

**Additional file 6: Table S3.** Analysis of *C*. *felis* proteins that did not cluster with other Holometabola.

<CLICK for link to Table S3>

**Additional file 7: Table S4.** Elements of the *C*. *felis* microbiome and associated *Wolbachia* phylogeny estimation.

<click for link to Table S4>

**Additional file 8: Table S5.** Coverage of corrected PacBio reads against all 16,622 polished assembly contigs.

<click for link to Table S5>

