## Supplemental Figures for "A chromosome-level assembly of the cat flea genome uncovers rampant gene duplication and genome size plasticity"

**Additional file 1: Fig. S1.** Assessing assembly fragmentation, gene duplication and repeat elements within the *C. felis* assembly. (**A**) Evaluating assembly fragmentation via mapping of scaffolds shorter than 1 Mb (n = 3,724) to scaffolds larger than 1 Mb (n = 9, "BIG9 scaffolds"). All but 2 short scaffolds mapped to a BIG9 scaffold at least once; confidence intervals are based on the probability of mapping to a single unique location. (**B**). Assessing the "genome completeness" of the *C. felis* full assembly and BIG9 scaffolds through comparison to eukaryote, arthropod and insect BUSCOs. (**C**) Tandem and proximal duplicate gene locations on BIG9 scaffold 1, (**D**) BIG9 scaffold 2, (**E**) BIG9 scaffold 3, (**F**) BIG9 scaffold 4, (**G**) BIG9 scaffold 5, (**H**) BIG9 scaffold 6, (**I**) BIG9 scaffold 7, (**J**) BIG9 scaffold 8, (**K**) BIG9 scaffold 9. (**L**) Duplications by proximity. Only true duplications (n=2) are shown. Red bars (*) depict "dispersed" clusters that span multiple scaffolds. (**M**) Dispersed duplicate gene locations across BIG9 scaffolds. (**N**) Distribution across BIG9 scaffolds of *C. felis* proteins annotated as "DNA integration" (GO:0015074, see **Additional file 2: Table S1** for specific accession numbers) and their relation to gene duplications. (**O**) Compilation of retroelements, DNA transposons and other repeat elements predicted across the BIG9 scaffolds. Overall totals are highlighted yellow. (**P**) tRNA gene abundances and (**Q**) codon usage/amino acid for select Holometabola.

A

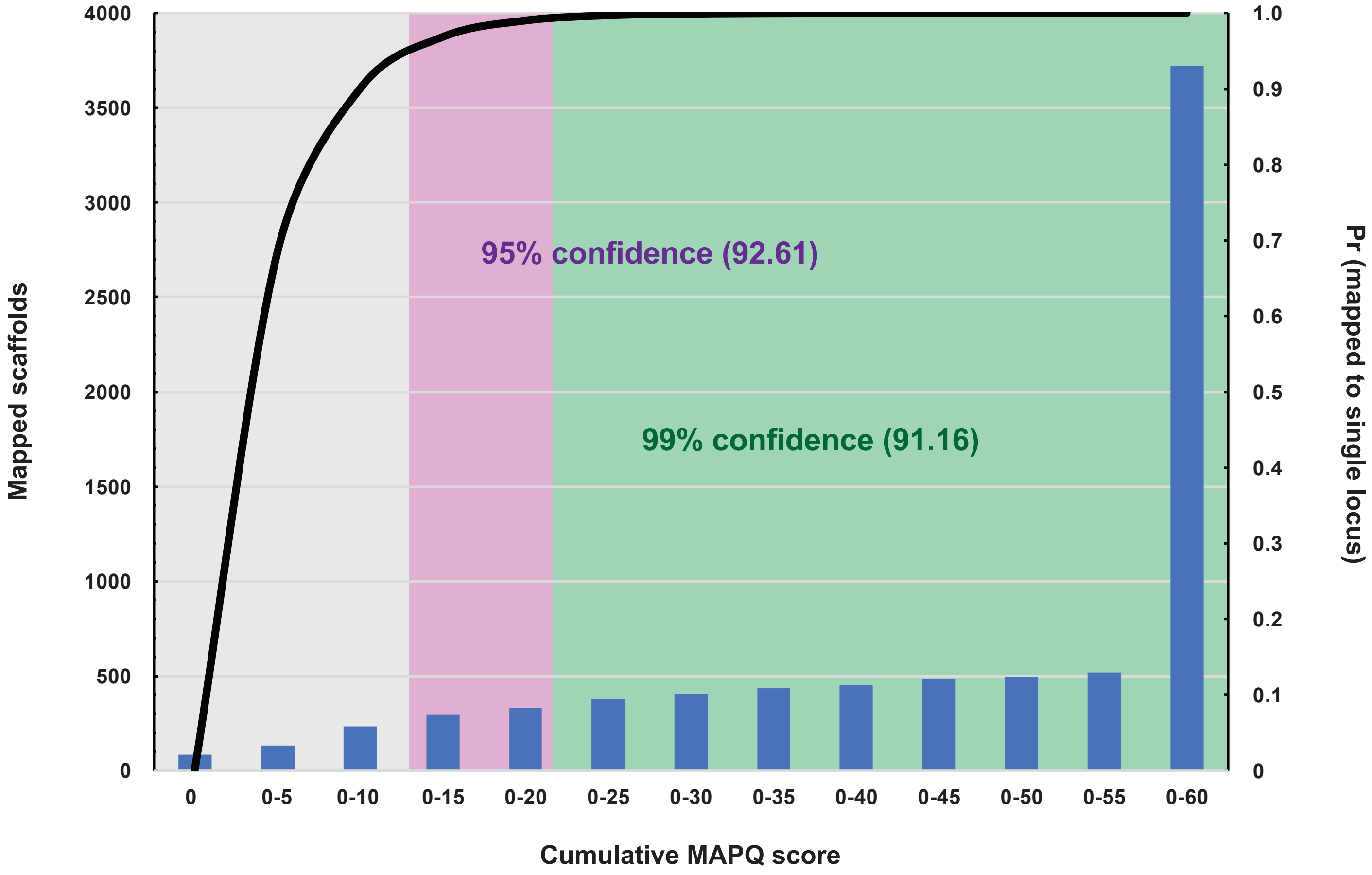

4000
3500
3000
2500
2000
1500
1000
500
0
1.0
0.9
0.8
0.7
0.6
0.5
0.4
0.3
0.2
0.1
0
Mapped scaffolds
Pr (mapped to single locus)
95% confidence (92.61)
99% confidence (91.16)
0
0-5
0-10
0-15
0-20
0-25
0-30
0-35
0-40
0-45
0-50
0-55
0-60
Cumulative MAPQ score


B

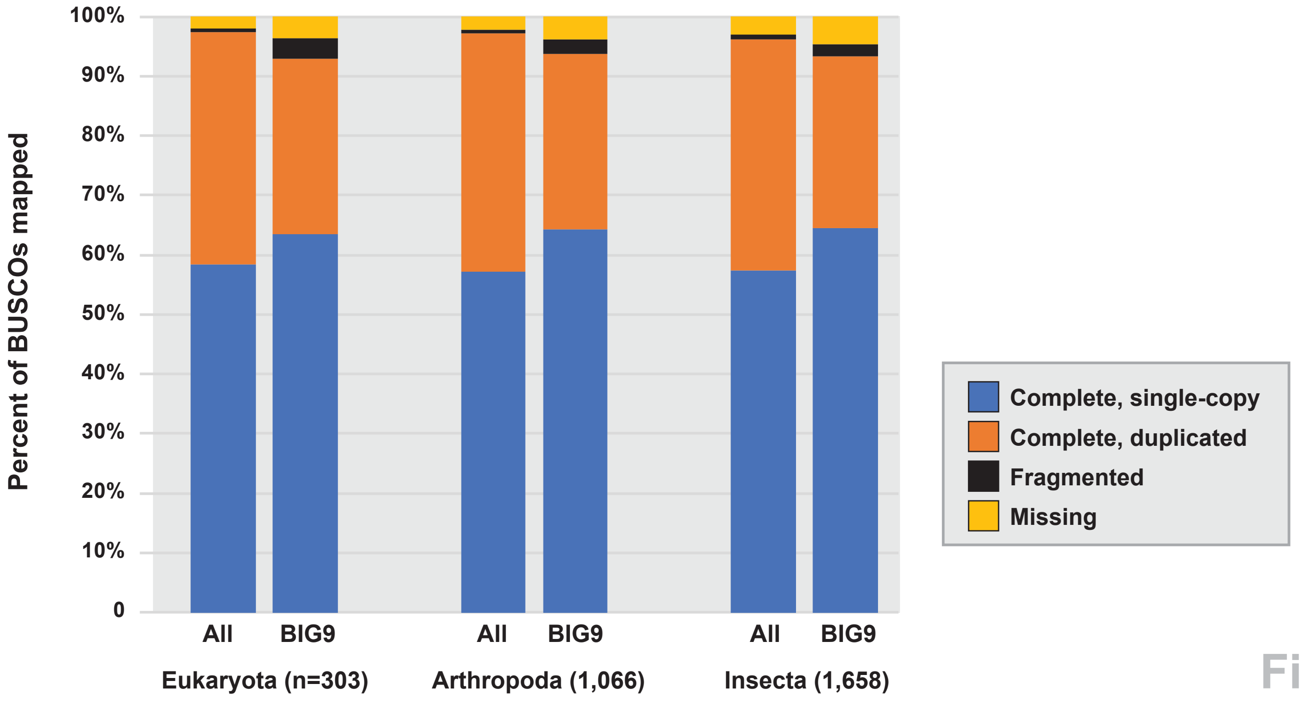

100%
90%
80%
70%
60%
50%
40%
30%
20%
10%
0
Percent of BUSCOs mapped
All
BIG9
All
BIG9
All
BIG9
Eukaryota (n=303)
Arthropoda (1,066)
Insecta (1,658)
Complete, single-copy
Complete, duplicated
Fragmented
Missing


Fig. S1

**C** **Scaffold 1 (NW_20538040.1)** **187,887,292 bp**

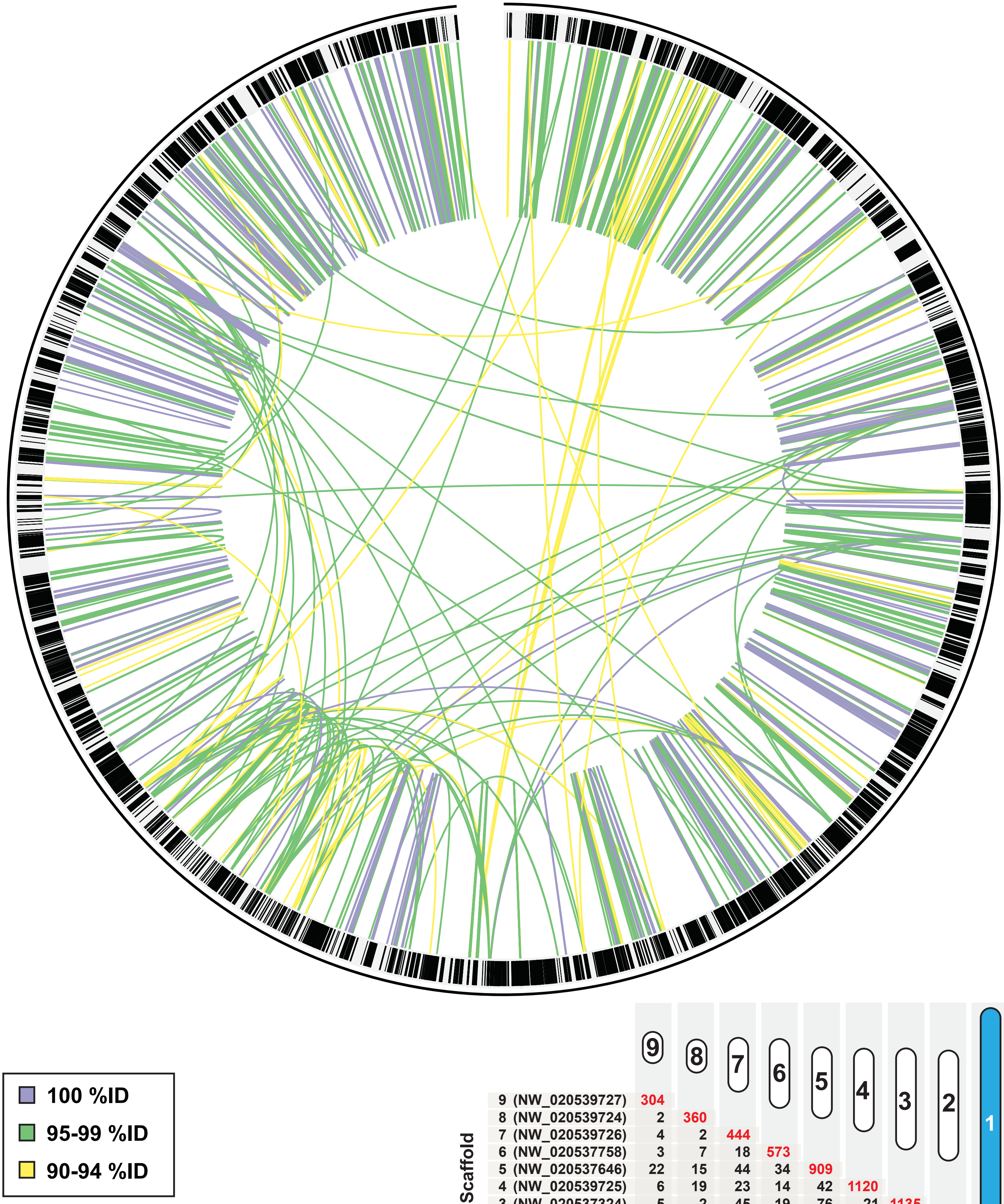


| Scaffold | 9 | 8 | 7 | 6 | 5 | 4 | 3 | 2 | 1 |
|---|---|---|---|---|---|---|---|---|---|
| 9 (NW_020539727) | 304 | | | | | | | | |
| 8 (NW_020539724) | 2 | 360 | | | | | | | |
| 7 (NW_020539726) | 4 | 2 | 444 | | | | | | |
| 6 (NW_020537758) | 3 | 7 | 18 | 573 | | | | | |
| 5 (NW_020537646) | 22 | 15 | 44 | 34 | 909 | | | | |
| 4 (NW_020539725) | 6 | 19 | 23 | 14 | 42 | 1120 | | | |
| 3 (NW_020537324) | 5 | 2 | 45 | 19 | 76 | 21 | 1135 | | |
| 2 (NW_020536999) | 4 | 13 | 48 | 32 | 64 | 43 | 68 | 1320 | |
| 1 (NW_020538040) | 23 | 5 | 91 | 47 | 137 | 52 | 81 | 137 | 1314 |

Fig. S1

D

Scaffold 2 (NW_20536999.1)

95,519,864 bp

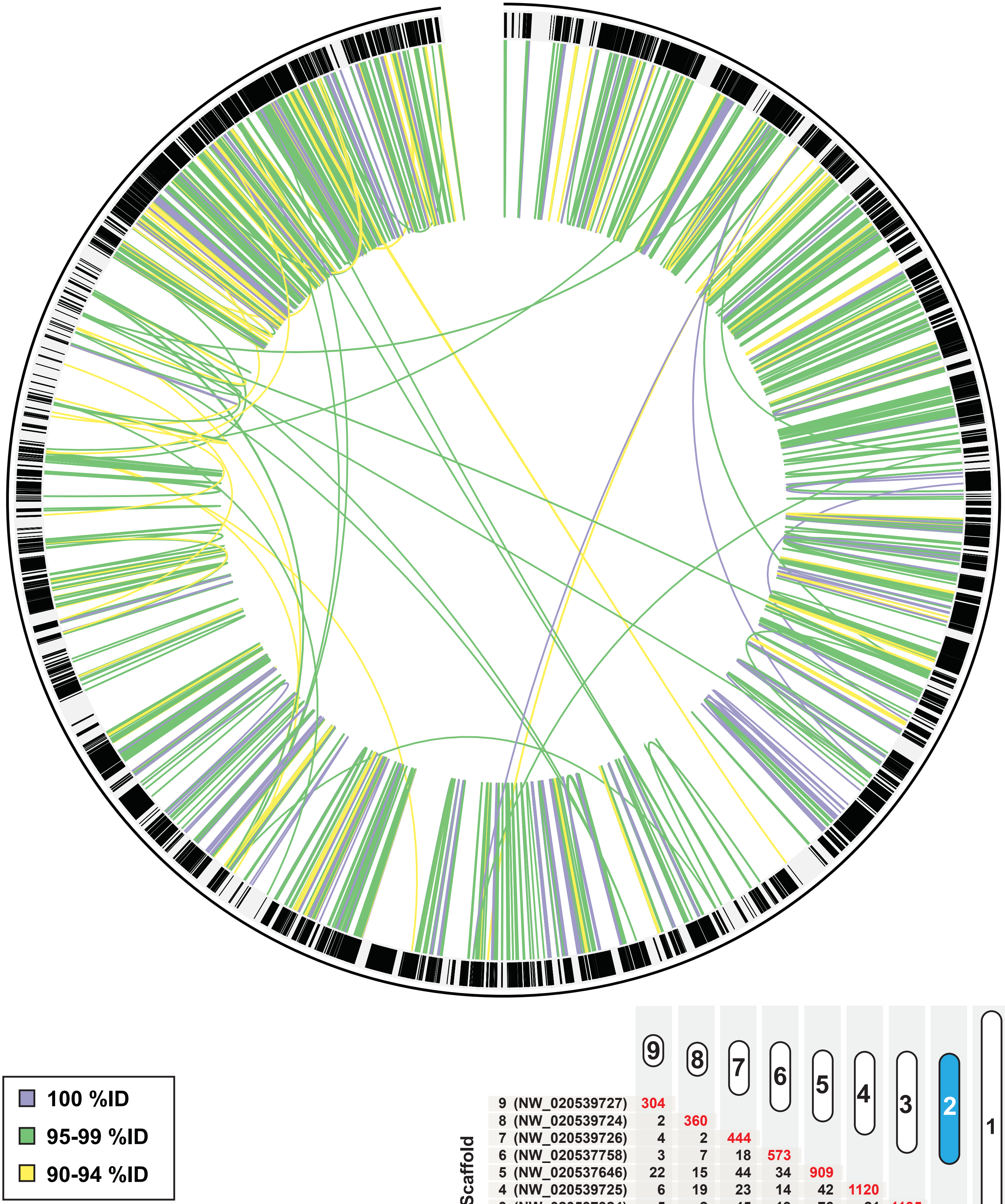


| Scaffold | 9 | 8 | 7 | 6 | 5 | 4 | 3 | 2 | 1 |
|---|---|---|---|---|---|---|---|---|---|
| 9 (NW_020539727) | 304 | | | | | | | | |
| 8 (NW_020539724) | 2 | 360 | | | | | | | |
| 7 (NW_020539726) | 4 | 2 | 444 | | | | | | |
| 6 (NW_020537758) | 3 | 7 | 18 | 573 | | | | | |
| 5 (NW_020537646) | 22 | 15 | 44 | 34 | 909 | | | | |
| 4 (NW_020539725) | 6 | 19 | 23 | 14 | 42 | 1120 | | | |
| 3 (NW_020537324) | 5 | 2 | 45 | 19 | 76 | 21 | 1135 | | |
| 2 (NW_020536999) | 4 | 13 | 48 | 32 | 64 | 43 | 68 | 1320 | |
| 1 (NW_020538040) | 23 | 5 | 91 | 47 | 137 | 52 | 81 | 137 | 1314 |

Fig. S1

**E** **Scaffold 3 (NW_20537324.1)** **87,719,082 bp**

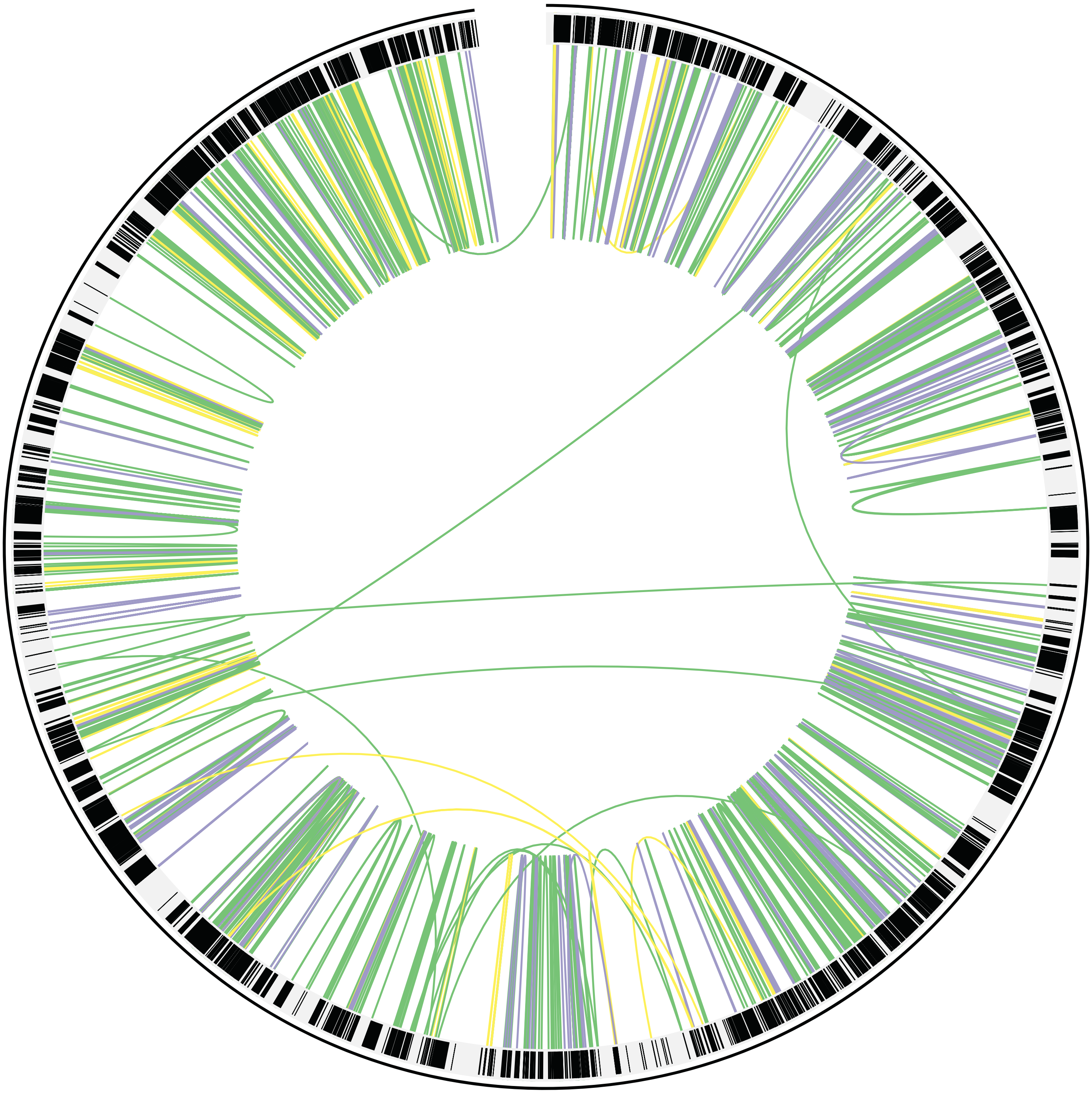

- 100 %ID
- 95-99 %ID
- 90-94 %ID

| Scaffold | 9 | 8 | 7 | 6 | 5 | 4 | 3 | 2 | 1 |
|---|---|---|---|---|---|---|---|---|---|
| 9 (NW_020539727) | 304 | | | | | | | | |
| 8 (NW_020539724) | 2 | 360 | | | | | | | |
| 7 (NW_020539726) | 4 | 2 | 444 | | | | | | |
| 6 (NW_020537758) | 3 | 7 | 18 | 573 | | | | | |
| 5 (NW_020537646) | 22 | 15 | 44 | 34 | 909 | | | | |
| 4 (NW_020539725) | 6 | 19 | 23 | 14 | 42 | 1120 | | | |
| 3 (NW_020537324) | 5 | 2 | 45 | 19 | 76 | 21 | 1135 | | |
| 2 (NW_020536999) | 4 | 13 | 48 | 32 | 64 | 43 | 68 | 1320 | |
| 1 (NW_020538040) | 23 | 5 | 91 | 47 | 137 | 52 | 81 | 137 | 1314 |

Fig. S1

**F** **Scaffold 4 (NW_020539725.1)** **71,713,785 bp**

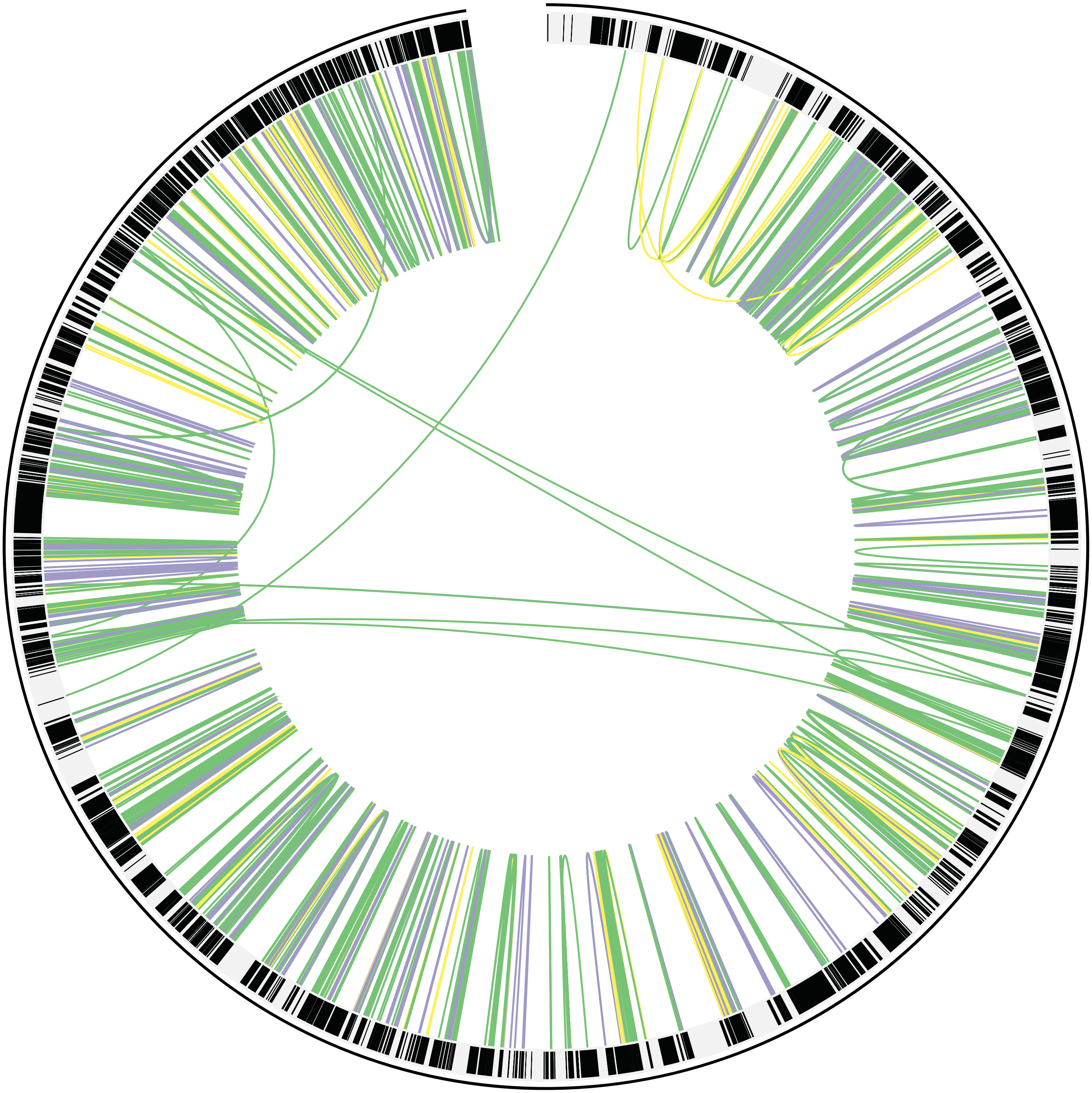

- 100 %ID
- 95-99 %ID
- 90-94 %ID

| Scaffold | 9 | 8 | 7 | 6 | 5 | 4 | 3 | 2 | 1 |
|---|---|---|---|---|---|---|---|---|---|
| 9 (NW_020539727) | 304 | | | | | | | | |
| 8 (NW_020539724) | 2 | 360 | | | | | | | |
| 7 (NW_020539726) | 4 | 2 | 444 | | | | | | |
| 6 (NW_020537758) | 3 | 7 | 18 | 573 | | | | | |
| 5 (NW_020537646) | 22 | 15 | 44 | 34 | 909 | | | | |
| 4 (NW_020539725) | 6 | 19 | 23 | 14 | 42 | 1120 | | | |
| 3 (NW_020537324) | 5 | 2 | 45 | 19 | 76 | 21 | 1135 | | |
| 2 (NW_020536999) | 4 | 13 | 48 | 32 | 64 | 43 | 68 | 1320 | |
| 1 (NW_020538040) | 23 | 5 | 91 | 47 | 137 | 52 | 81 | 137 | 1314 |

Fig. S1

**G** **Scaffold 5 (NW_020537646.1)** **61,547,676 bp**

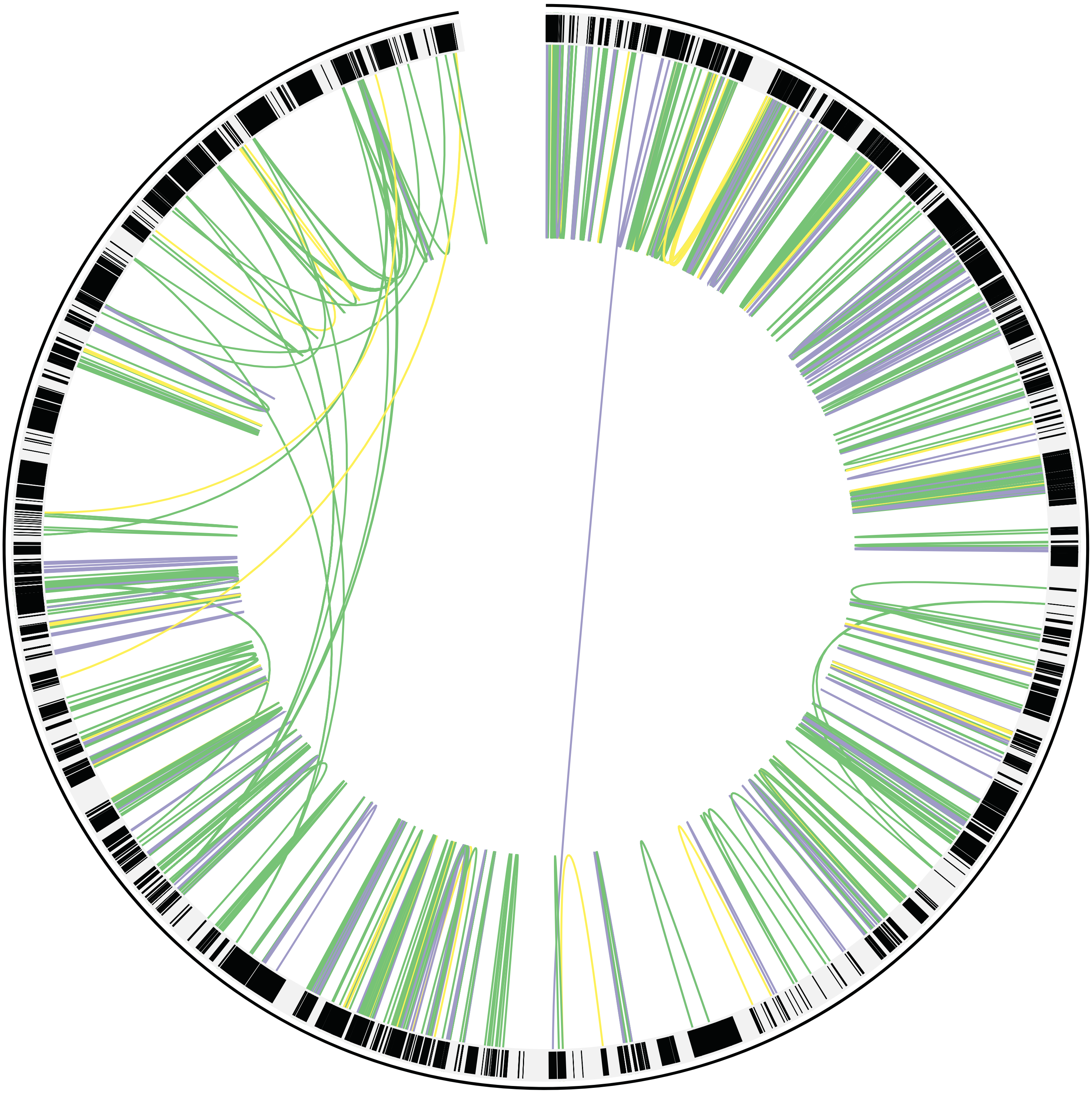

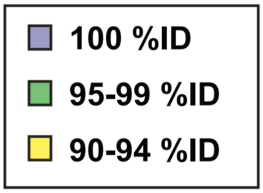


| Scaffold | 9 | 8 | 7 | 6 | 5 | 4 | 3 | 2 | 1 |
|---|---|---|---|---|---|---|---|---|---|
| 9 (NW_020539727) | 304 | | | | | | | | |
| 8 (NW_020539724) | 2 | 360 | | | | | | | |
| 7 (NW_020539726) | 4 | 2 | 444 | | | | | | |
| 6 (NW_020537758) | 3 | 7 | 18 | 573 | | | | | |
| 5 (NW_020537646) | 22 | 15 | 44 | 34 | 909 | | | | |
| 4 (NW_020539725) | 6 | 19 | 23 | 14 | 42 | 1120 | | | |
| 3 (NW_020537324) | 5 | 2 | 45 | 19 | 76 | 21 | 1135 | | |
| 2 (NW_020536999) | 4 | 13 | 48 | 32 | 64 | 43 | 68 | 1320 | |
| 1 (NW_020538040) | 23 | 5 | 91 | 47 | 137 | 52 | 81 | 137 | 1314 |

Fig. S1

**H** **Scaffold 6 (NW_020537758.1)** **59,836,375 bp**

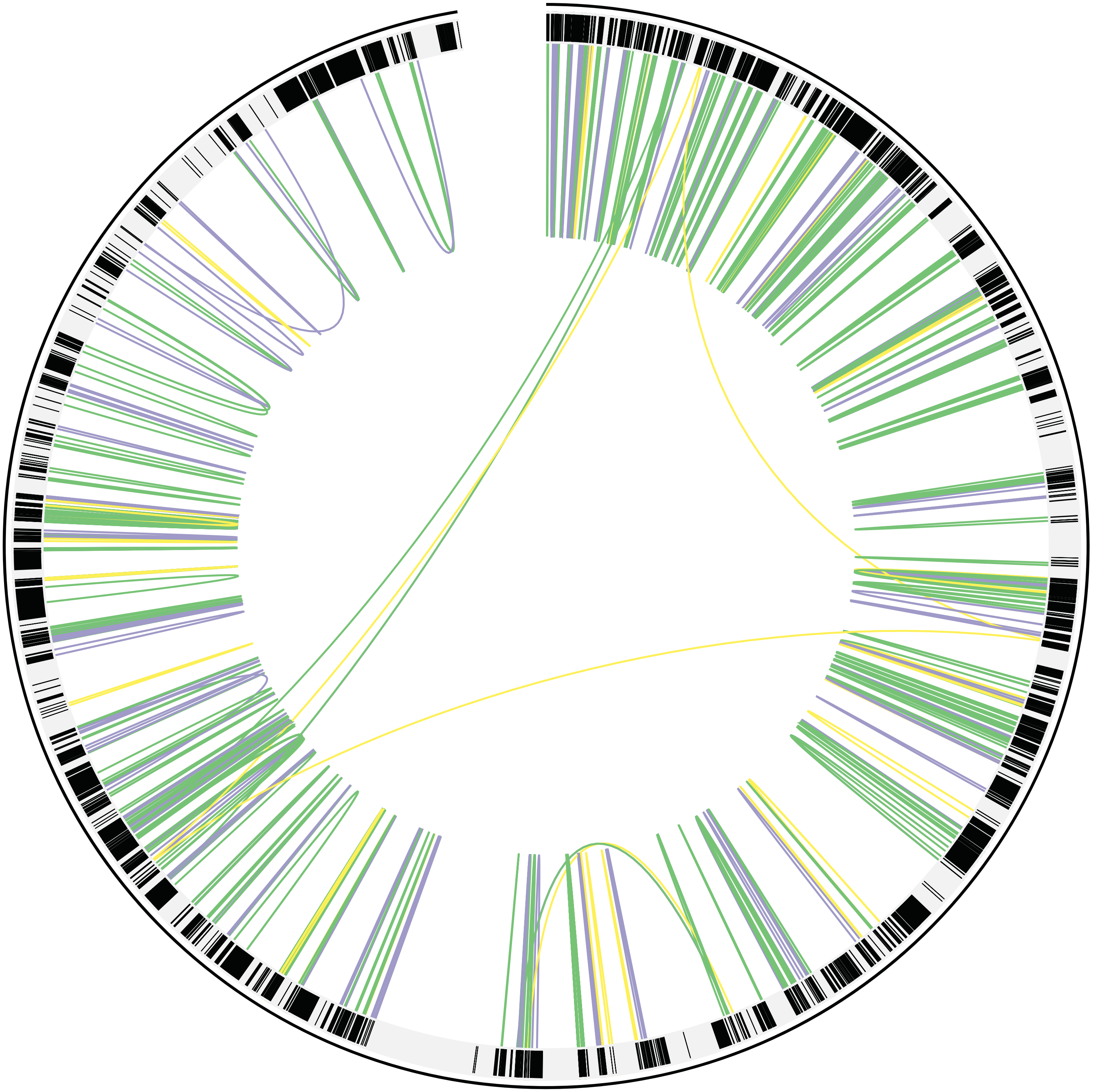

| Scaffold | 9 | 8 | 7 | 6 | 5 | 4 | 3 | 2 | 1 |
|---|---|---|---|---|---|---|---|---|---|
| 9 (NW_020539727) | 304 | | | | | | | | |
| 8 (NW_020539724) | 2 | 360 | | | | | | | |
| 7 (NW_020539726) | 4 | 2 | 444 | | | | | | |
| 6 (NW_020537758) | 3 | 7 | 18 | 573 | | | | | |
| 5 (NW_020537646) | 22 | 15 | 44 | 34 | 909 | | | | |
| 4 (NW_020539725) | 6 | 19 | 23 | 14 | 42 | 1120 | | | |
| 3 (NW_020537324) | 5 | 2 | 45 | 19 | 76 | 21 | 1135 | | |
| 2 (NW_020536999) | 4 | 13 | 48 | 32 | 64 | 43 | 68 | 1320 | |
| 1 (NW_020538040) | 23 | 5 | 91 | 47 | 137 | 52 | 81 | 137 | 1314 |

- 100 %ID
- 95-99 %ID
- 90-94 %ID

I

**Scaffold 7 (NW_020539726.1)** **46,735,595 bp**

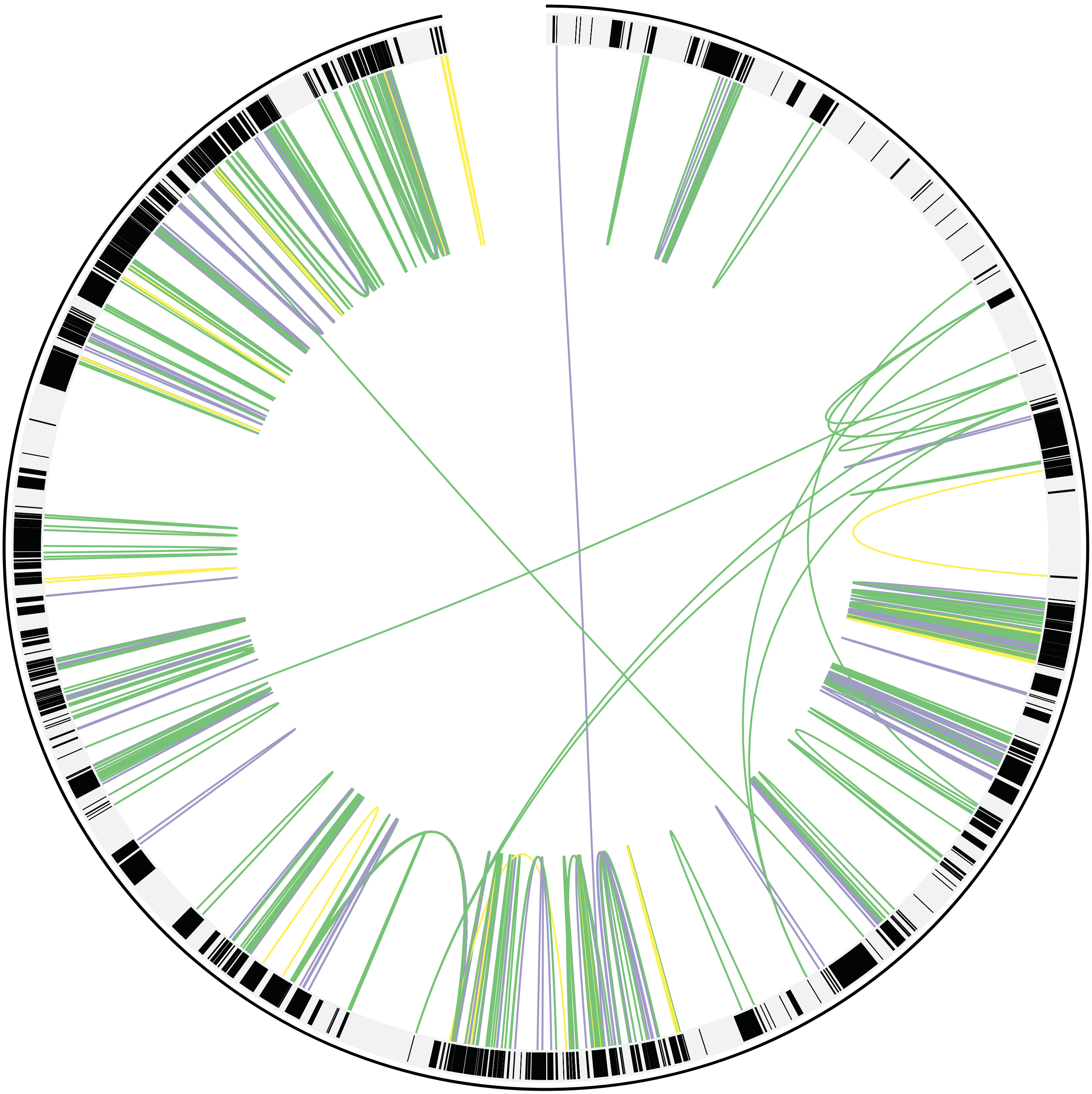

- 100 %ID
- 95-99 %ID
- 90-94 %ID

| Scaffold | 9 | 8 | 7 | 6 | 5 | 4 | 3 | 2 | 1 |
|---|---|---|---|---|---|---|---|---|---|
| 9 (NW_020539727) | 304 | | | | | | | | |
| 8 (NW_020539724) | 2 | 360 | | | | | | | |
| 7 (NW_020539726) | 4 | 2 | 444 | | | | | | |
| 6 (NW_020537758) | 3 | 7 | 18 | 573 | | | | | |
| 5 (NW_020537646) | 22 | 15 | 44 | 34 | 909 | | | | |
| 4 (NW_020539725) | 6 | 19 | 23 | 14 | 42 | 1120 | | | |
| 3 (NW_020537324) | 5 | 2 | 45 | 19 | 76 | 21 | 1135 | | |
| 2 (NW_020536999) | 4 | 13 | 48 | 32 | 64 | 43 | 68 | 1320 | |
| 1 (NW_020538040) | 23 | 5 | 91 | 47 | 137 | 52 | 81 | 137 | 1314 |

Fig. S1

**J** **Scaffold 8 (NW_020539724.1)** **28,733,183 bp**

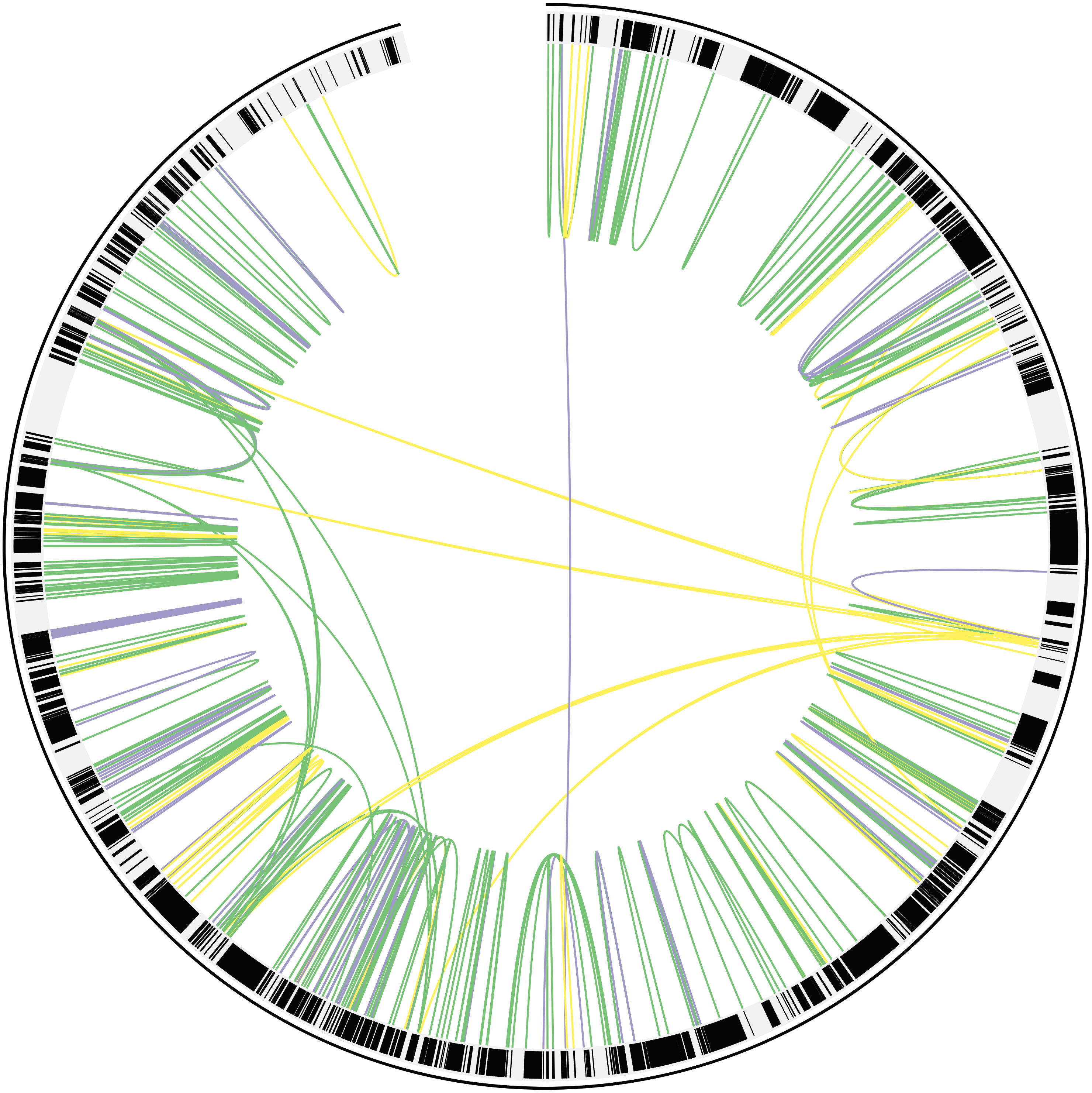

- 100 %ID
- 95-99 %ID
- 90-94 %ID

| Scaffold | 9 | 8 | 7 | 6 | 5 | 4 | 3 | 2 | 1 |
|---|---|---|---|---|---|---|---|---|---|
| 9 (NW_020539727) | 304 | | | | | | | | |
| 8 (NW_020539724) | 2 | 360 | | | | | | | |
| 7 (NW_020539726) | 4 | 2 | 444 | | | | | | |
| 6 (NW_020537758) | 3 | 7 | 18 | 573 | | | | | |
| 5 (NW_020537646) | 22 | 15 | 44 | 34 | 909 | | | | |
| 4 (NW_020539725) | 6 | 19 | 23 | 14 | 42 | 1120 | | | |
| 3 (NW_020537324) | 5 | 2 | 45 | 19 | 76 | 21 | 1135 | | |
| 2 (NW_020536999) | 4 | 13 | 48 | 32 | 64 | 43 | 68 | 1320 | |
| 1 (NW_020538040) | 23 | 5 | 91 | 47 | 137 | 52 | 81 | 137 | 1314 |

Fig. S1

**K** **Scaffold 9 (NW_020539727.1)** **25,564,690 bp**

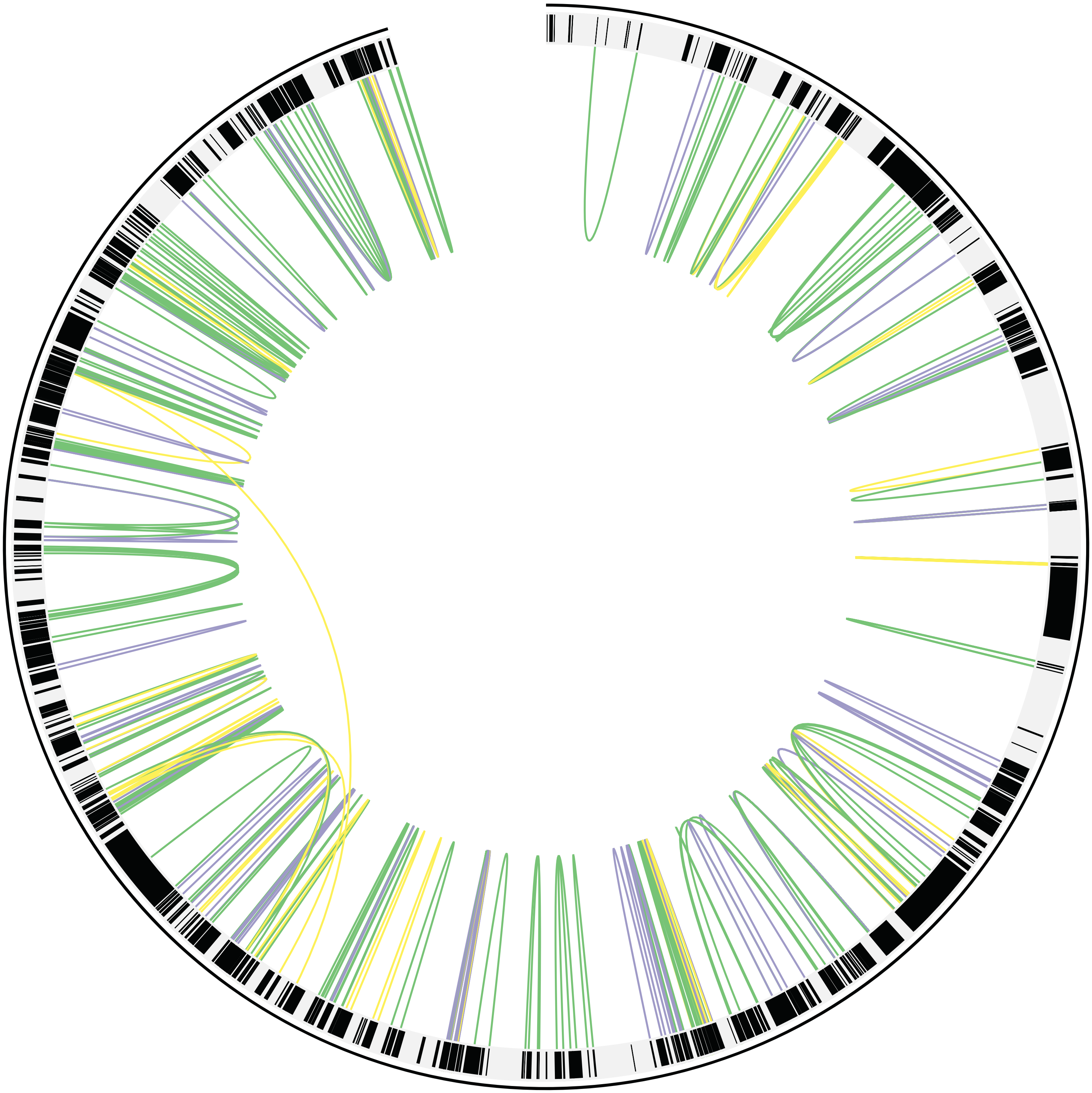

- 100 %ID
- 95-99 %ID
- 90-94 %ID

| Scaffold | 9 | 8 | 7 | 6 | 5 | 4 | 3 | 2 | 1 |
|---|---|---|---|---|---|---|---|---|---|
| 9 (NW_020539727) | 304 | | | | | | | | |
| 8 (NW_020539724) | 2 | 360 | | | | | | | |
| 7 (NW_020539726) | 4 | 2 | 444 | | | | | | |
| 6 (NW_020537758) | 3 | 7 | 18 | 573 | | | | | |
| 5 (NW_020537646) | 22 | 15 | 44 | 34 | 909 | | | | |
| 4 (NW_020539725) | 6 | 19 | 23 | 14 | 42 | 1120 | | | |
| 3 (NW_020537324) | 5 | 2 | 45 | 19 | 76 | 21 | 1135 | | |
| 2 (NW_020536999) | 4 | 13 | 48 | 32 | 64 | 43 | 68 | 1320 | |
| 1 (NW_020538040) | 23 | 5 | 91 | 47 | 137 | 52 | 81 | 137 | 1314 |

Fig. S1

L

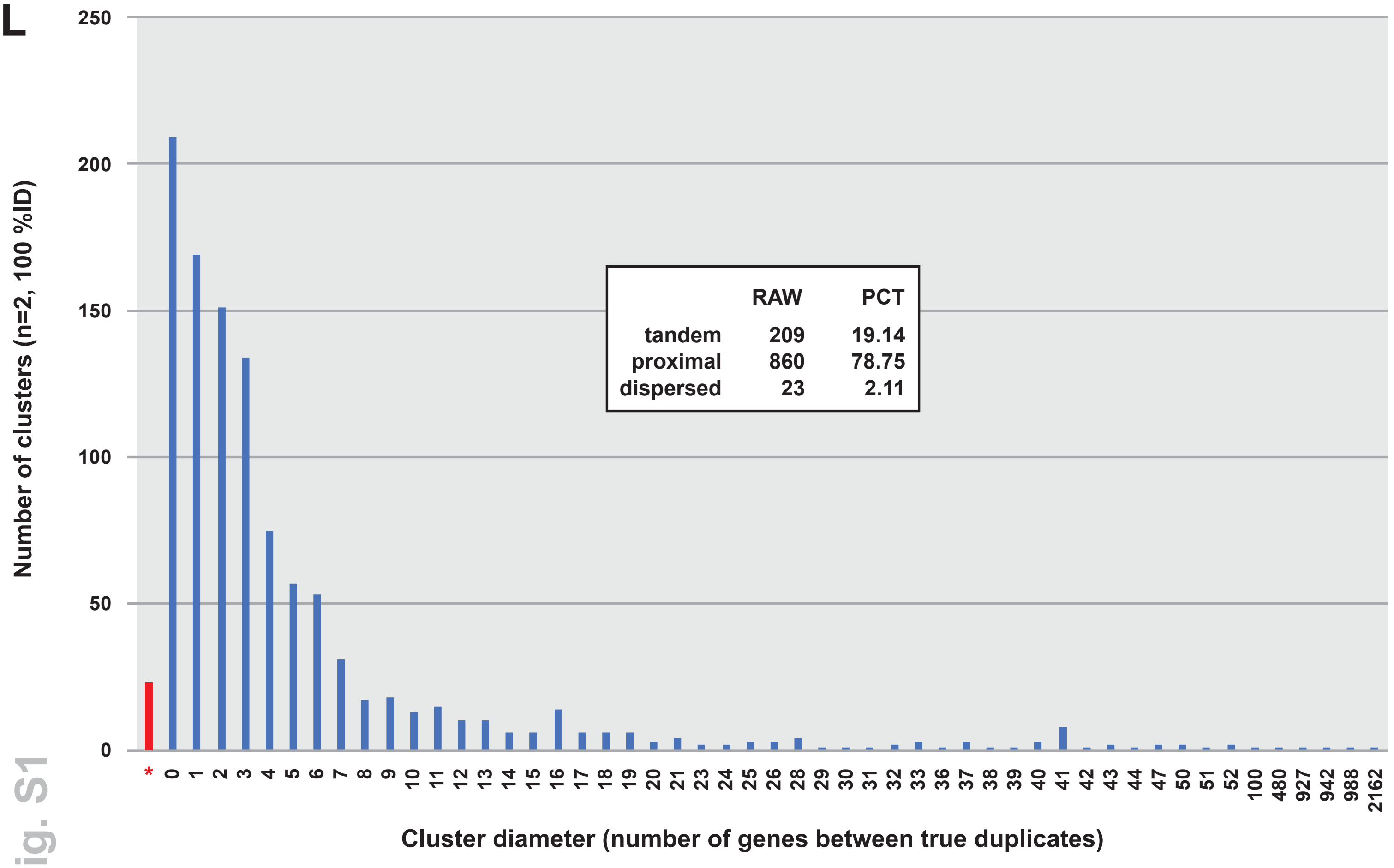


| | RAW | PCT |
|---|---|---|
| tandem | 209 | 19.14 |
| proximal | 860 | 78.75 |
| dispersed | 23 | 2.11 |

Fig. S1

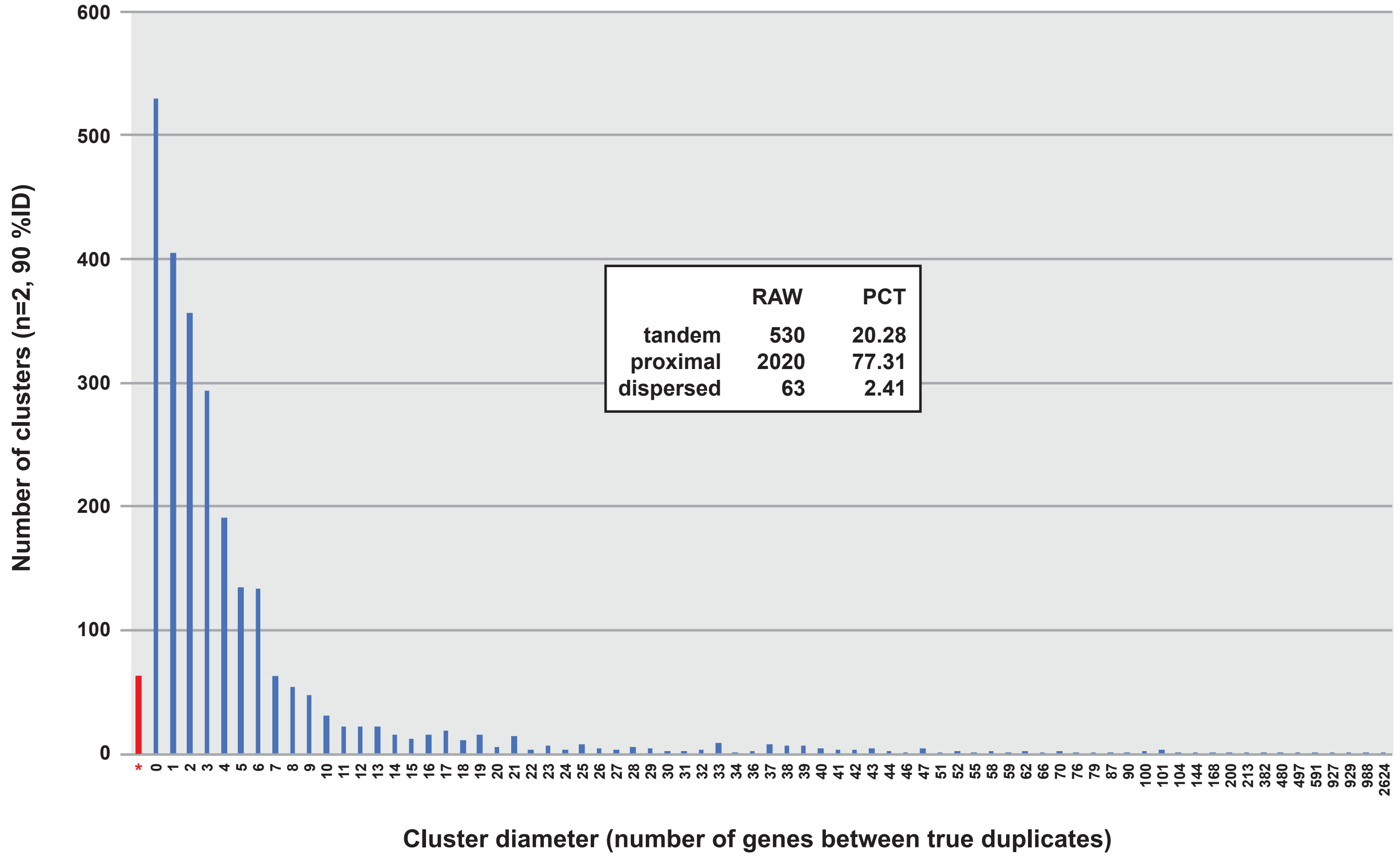


| | RAW | PCT |
|---|---|---|
| tandem | 530 | 20.28 |
| proximal | 2020 | 77.31 |
| dispersed | 63 | 2.41 |

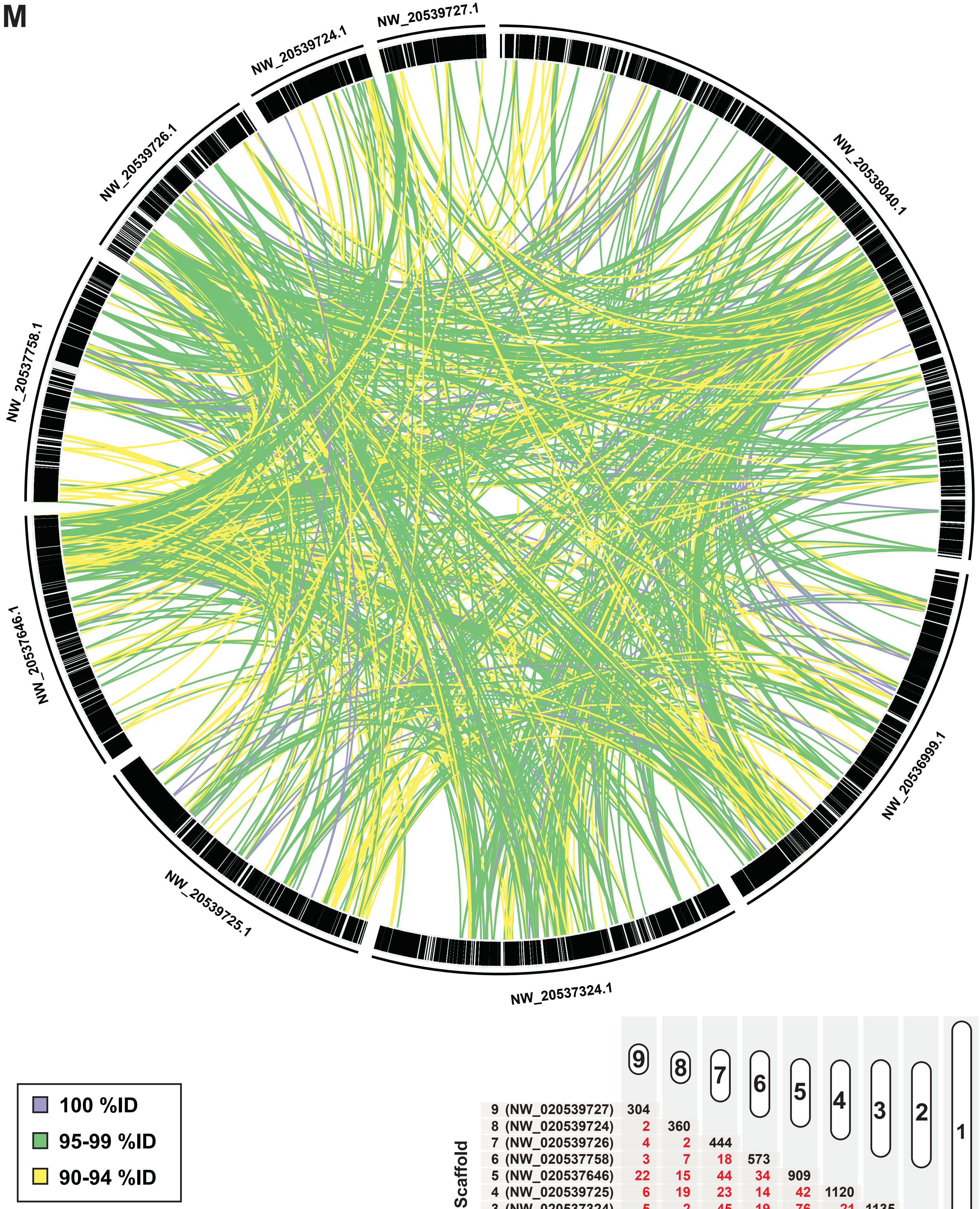


| Scaffold | 9 | 8 | 7 | 6 | 5 | 4 | 3 | 2 | 1 |
|---|---|---|---|---|---|---|---|---|---|
| 9 (NW_020539727) | 304 | | | | | | | | |
| 8 (NW_020539724) | 2 | 360 | | | | | | | |
| 7 (NW_020539726) | 4 | 2 | 444 | | | | | | |
| 6 (NW_020537758) | 3 | 7 | 18 | 573 | | | | | |
| 5 (NW_020537646) | 22 | 15 | 44 | 34 | 909 | | | | |
| 4 (NW_020539725) | 6 | 19 | 23 | 14 | 42 | 1120 | | | |
| 3 (NW_020537324) | 5 | 2 | 45 | 19 | 76 | 21 | 1135 | | |
| 2 (NW_020536999) | 4 | 13 | 48 | 32 | 64 | 43 | 68 | 1320 | |
| 1 (NW_020538040) | 23 | 5 | 91 | 47 | 137 | 52 | 81 | 137 | 1314 |

Fig. S1

N

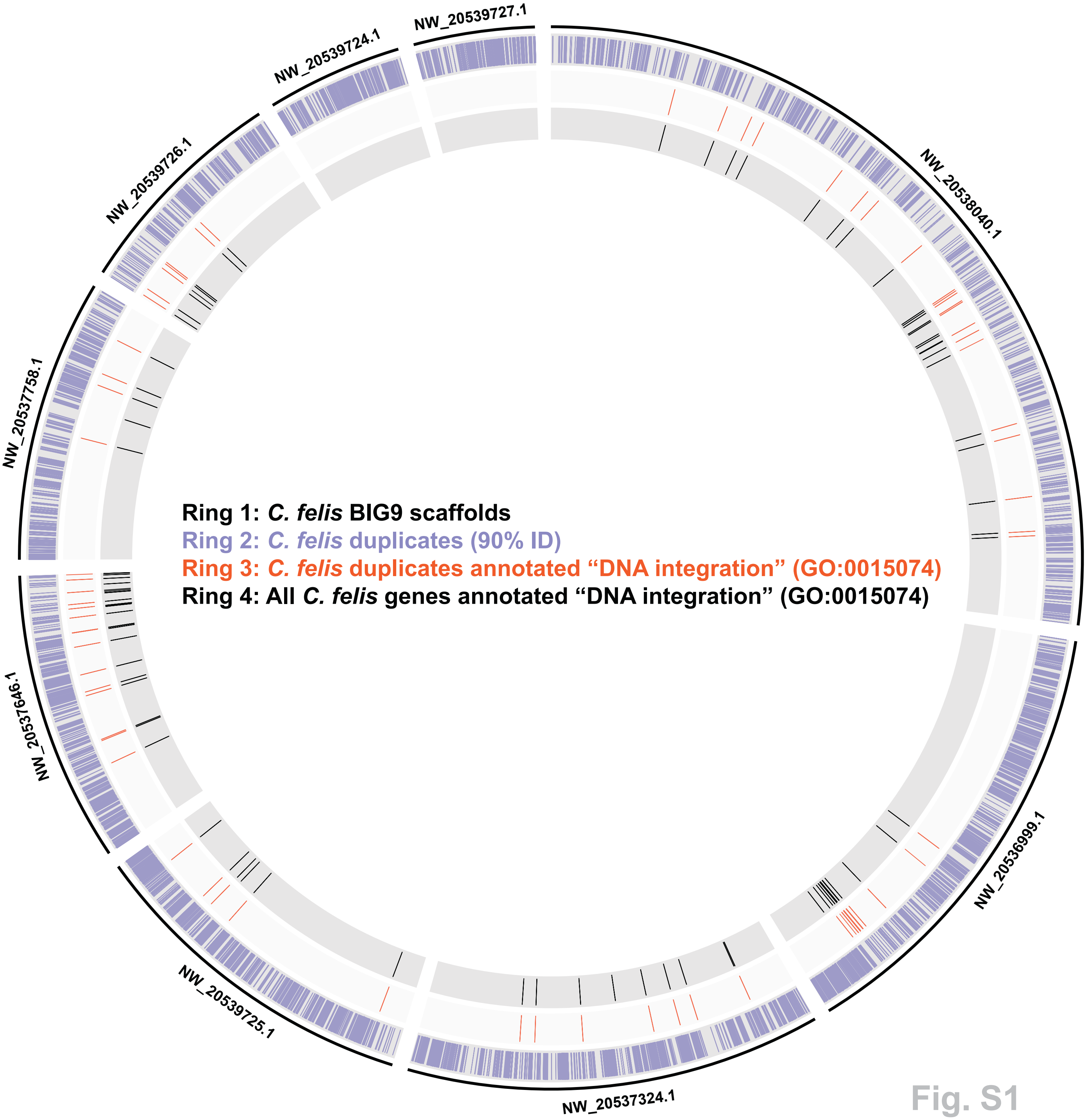
NW_20539727.1
NW_20539724.1
NW_20539726.1
NW_20537758.1
NW_20537646.1
NW_20539725.1
NW_20537324.1
NW_20536999.1
NW_20538040.1
Ring 1: *C. felis* BIG9 scaffolds
Ring 2: *C. felis* duplicates (90% ID)
Ring 3: *C. felis* duplicates annotated "DNA integration" (GO:0015074)
Ring 4: All *C. felis* genes annotated "DNA integration" (GO:0015074)

Fig. S1

O

| Superfamily | Total no. | Total Length | % of BIG9 |
|---|---|---|---|
| Retroelements | 61,071 | 12,619,486 | 1.9295 |
| SINEs | 336 | 26,535 | 0.0041 |
| LINEs | 34,965 | 8,515,826 | 1.302 |
| Penelope | 3,086 | 499,380 | 0.0764 |
| CRE/SLACS | 2 | 162 | 0.0000 |
| L2/CR1/Rex | 10,015 | 1,081,344 | 0.1653 |
| R2/R4/NeSL | 491 | 177,375 | 0.0271 |
| L1/CIN4 | 445 | 61,750 | 0.0094 |
| RTEs | 713 | 133,953 | 0.0205 |
| Other LINEs | 20,213 | 6,561,862 | 1.0033 |
| LTRs | 25,770 | 4,077,125 | 0.6234 |
| Unclassifed LTR | 94 | 5,062 | 0.0008 |
| Bel/Pao | 5,675 | 989,527 | 0.1513 |
| Ty1/Copia | 991 | 137,555 | 0.0210 |
| Gypsy/DIRS1 | 19,010 | 2,944,981 | 0.4503 |
| DNA transposons | 182,844 | 24,537,653 | 3.752 |
| Unclassified DNA transposons | 91,977 | 12,373,802 | 1.8919 |
| hAT | 9,939 | 960,540 | 0.1469 |
| IS630-Tc1-Mariner | 36,131 | 6,148,685 | 0.9401 |
| En-Spm | 80 | 5,537 | 0.0008 |
| MuDR | 3,801 | 352,443 | 0.0539 |
| PiggyBac | 1,934 | 302,651 | 0.0463 |
| Tourist/Harbinger/PIF | 1,935 | 156,040 | 0.0239 |
| Zator | 15,858 | 1,770,968 | 0.2708 |
| Sola | 408 | 62,065 | 0.0095 |
| Other DNA transposons | 20,781 | 2,404,922 | 0.3677 |
| Rolling Circles | 29,853 | 4,381,462 | 0.6699 |
| Unclassified Repeats | 1,866 | 232,894 | 0.0356 |
| rRNA/tRNA | 3,703 | 809,578 | 0.1238 |
| Satellites | 206 | 34,766 | 0.0053 |
| Simple Repeats | 42,5231 | 19,488,965 | 2.9798 |
| Low Complexity | 62,484 | 3,137,132 | 0.4797 |
| Total Repeat Elements: | 767,258 | 65,241,936 | 9.975 |

Fig. S1

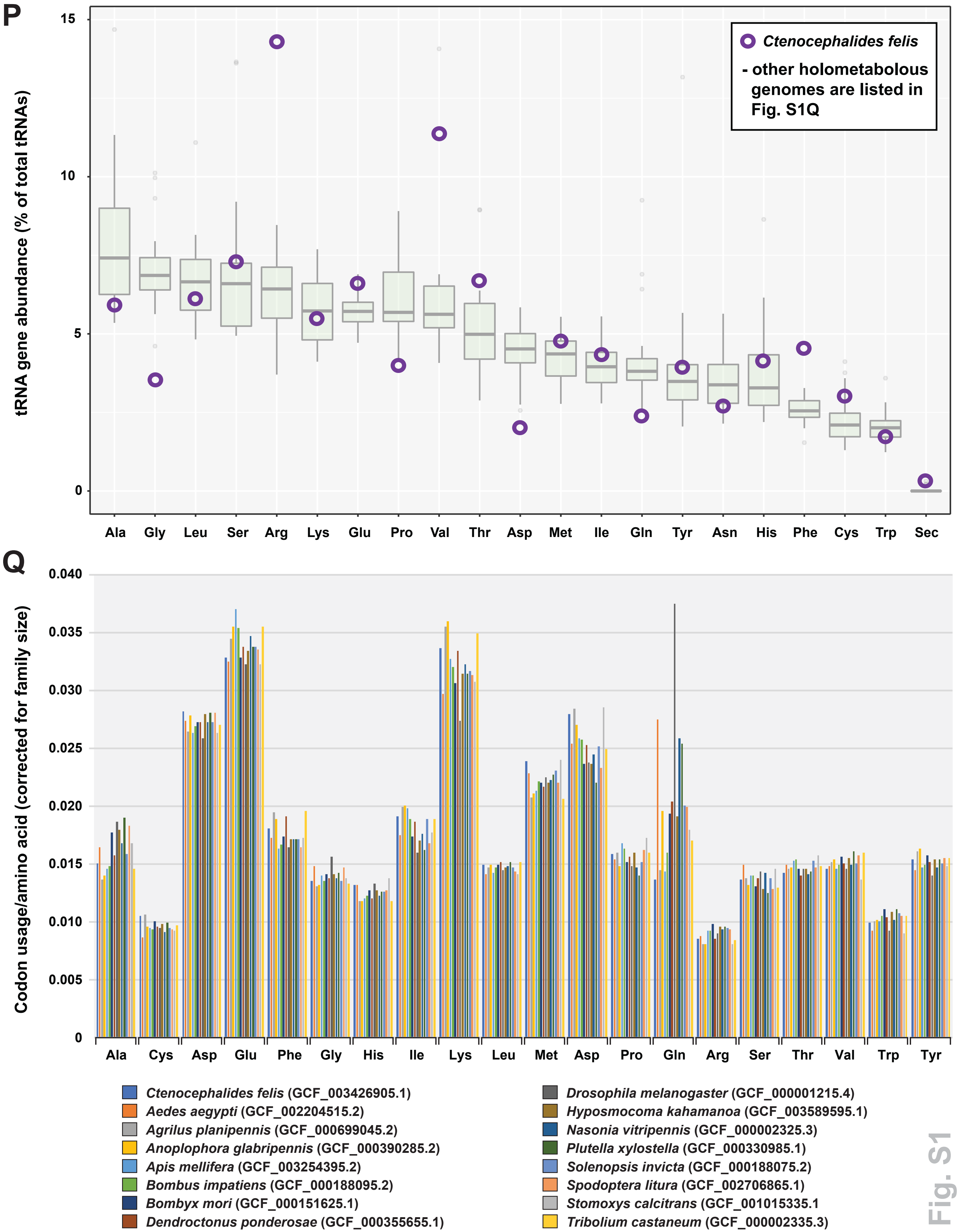
P
Ctenocephalides felis
- other holometabolous genomes are listed in Fig. S1Q
tRNA gene abundance (% of total tRNAs)
15
10
5
0
Ala Gly Leu Ser Arg Lys Glu Pro Val Thr Asp Met Ile Gln Tyr Asn His Phe Cys Trp Sec
Q
Codon usage/amino acid (corrected for family size)
0.040
0.035
0.030
0.025
0.020
0.015
0.010
0.005
0
Ala Cys Asp Glu Phe Gly His Ile Lys Leu Met Asp Pro Gln Arg Ser Thr Val Trp Tyr
Ctenocephalides felis (GCF_003426905.1)
Aedes aegypti (GCF_002204515.2)
Agrilus planipennis (GCF_000699045.2)
Anoplophora glabripennis (GCF_000390285.2)
Apis mellifera (GCF_003254395.2)
Bombus impatiens (GCF_000188095.2)
Bombyx mori (GCF_000151625.1)
Dendroctonus ponderosae (GCF_000355655.1)
Drosophila melanogaster (GCF_000001215.4)
Hyposmocoma kahamanoa (GCF_003589595.1)
Nasonia vitripennis (GCF_000002325.3)
Plutella xylostella (GCF_000330985.1)
Solenopsis invicta (GCF_000188075.2)
Spodoptera litura (GCF_002706865.1)
Stomoxys calcitrans (GCF_001015335.1
Tribolium castaneum (GCF_000002335.3)

Fig. S1

**Additional file 2: Table S1.** Functional predictions and enrichment analysis of *C. felis* proteins.

<click for link to Table S1>

**Additional file 3: Fig. S2.** Representative histograms produced by flow cytometry showing the peak positions of the 2C nuclei of *Drosophila melanogaster* (left) and *D. virilis* (center) female standards, and individual *C. felis* females (right) from the sequenced EL strain. (**A**) A 434 Mb flea. (**B**) A 553 Mb flea. All peaks have CV < 1.5 and > 500 nuclei under the statistical gates (red lines spanning the 2C peaks).

A

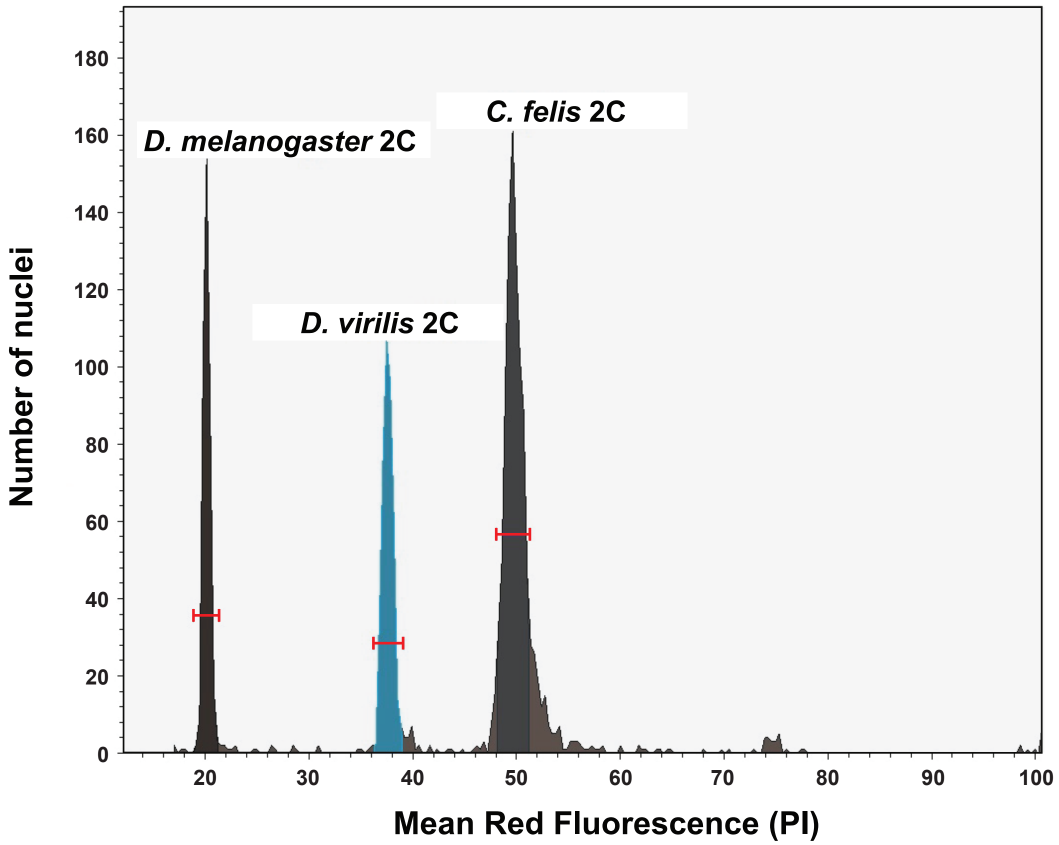

D. melanogaster 2C
C. felis 2C
D. virilis 2C
Number of nuclei
Mean Red Fluorescence (PI)
0
20
40
60
80
100
120
140
160
180
20
30
40
50
60
70
80
90
100


B

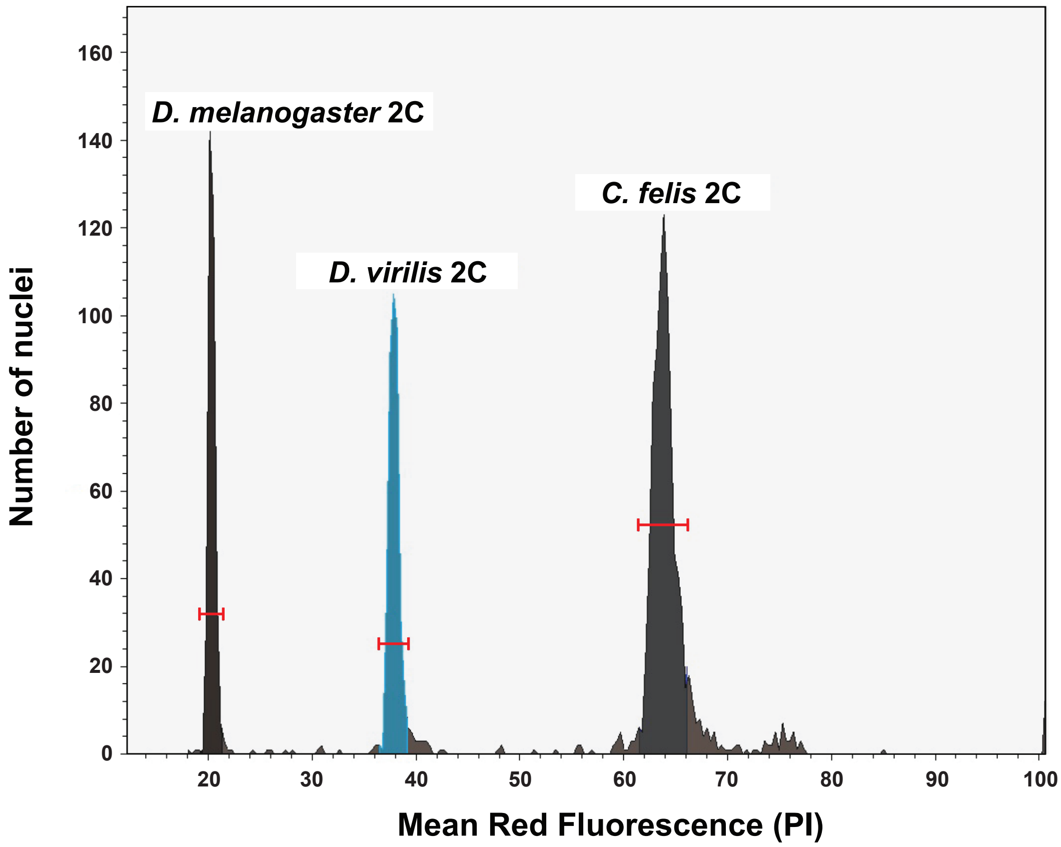

D. melanogaster 2C
C. felis 2C
D. virilis 2C
Number of nuclei
Mean Red Fluorescence (PI)
0
20
40
60
80
100
120
140
160
20
30
40
50
60
70
80
90
100


Fig. S2

**Additional file 4: Fig. S3.** Phylogenomics analysis of select Holometabola. (**A**) Assessment of holometabolan accessory genomes. (**B**) *Top:* Identification of conserved protein families present in select taxa from each holometabolan order but absent from *C. felis*. *Bottom:* Protein families conserved across all sequenced holometabolan genomes except *C. felis* (see **Additional file 5: Table S2**). Four assemblies were identified as particularly patchy (*Oryctes borbonicus*, *Operophtera brumata*, *Heliothis virescens*, and *Plutella xylostella*) and 100% conservation ("perfect") was also relaxed to exclude these taxa. Inset, redrawn phylogeny estimation of Holometabola [2].

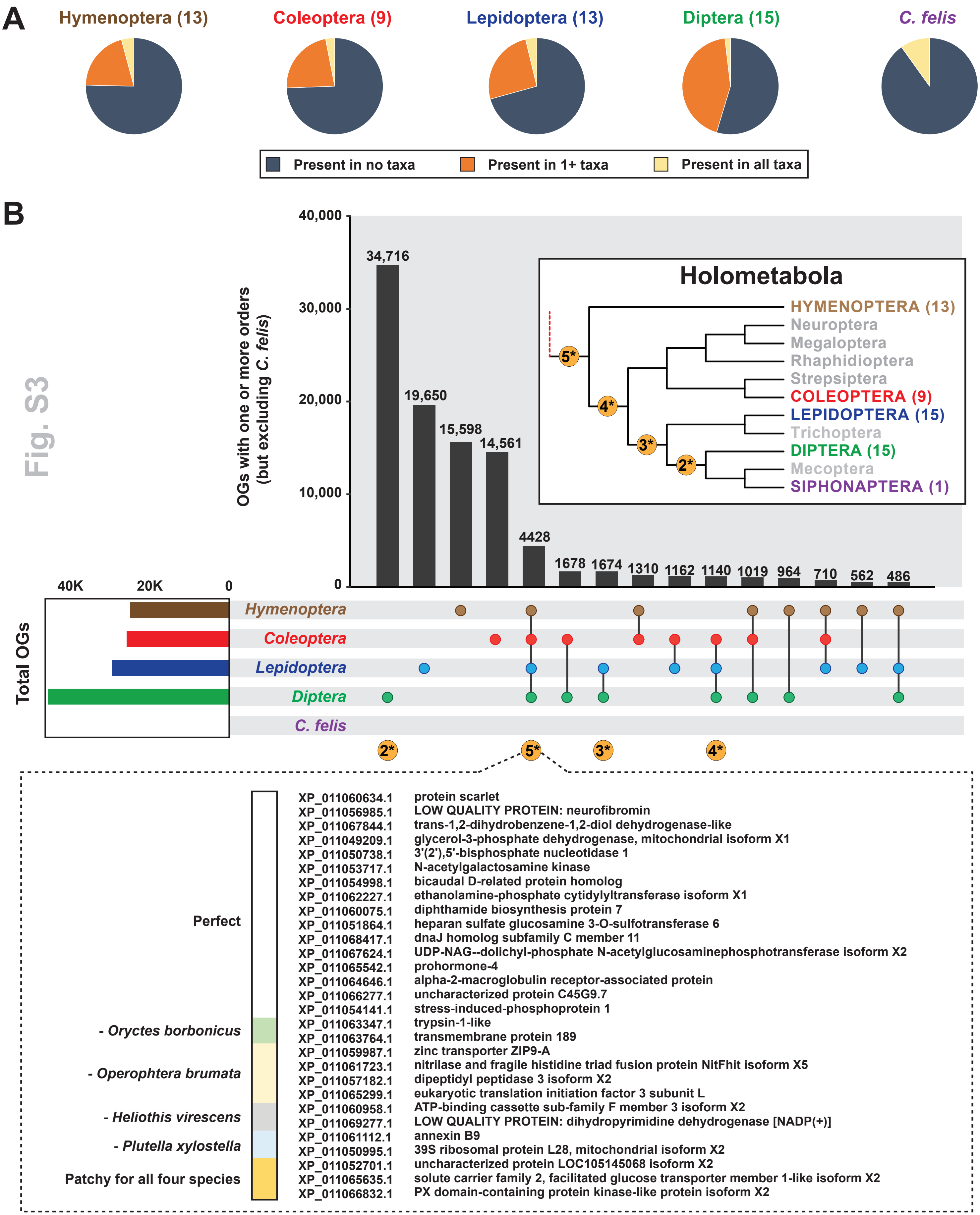
A
Hymenoptera (13)
Coleoptera (9)
Lepidoptera (13)
Diptera (15)
C. felis
Present in no taxa
Present in 1+ taxa
Present in all taxa
B
OGs with one or more orders
(but excluding C. felis)
40,000
30,000
20,000
10,000
0
34,716
19,650
15,598
14,561
4428
1678
1674
1310
1162
1140
1019
964
710
562
486
Holometabola
HYMENOPTERA (13)
Neuroptera
Megaloptera
Rhaphidioptera
Strepsiptera
COLEOPTERA (9)
LEPIDOPTERA (15)
Trichoptera
DIPTERA (15)
Mecoptera
SIPHONAPTERA (1)
5*
4*
3*
2*
Total OGs
40K
20K
0
Hymenoptera
Coleoptera
Lepidoptera
Diptera
C. felis
2*
5*
3*
4*
Perfect
- Oryctes borbonicus
- Operophtera brumata
- Heliothis virescens
- Plutella xylostella
Patchy for all four species
XP_011060634.1 protein scarlet
XP_011056985.1 LOW QUALITY PROTEIN: neurofibromin
XP_011067844.1 trans-1,2-dihydrobenzene-1,2-diol dehydrogenase-like
XP_011049209.1 glycerol-3-phosphate dehydrogenase, mitochondrial isoform X1
XP_011050738.1 3'(2'),5'-bisphosphate nucleotidase 1
XP_011053717.1 N-acetylgalactosamine kinase
XP_011054998.1 bicaudal D-related protein homolog
XP_011062227.1 ethanolamine-phosphate cytidylyltransferase isoform X1
XP_011060075.1 diphthamide biosynthesis protein 7
XP_011051864.1 heparan sulfate glucosamine 3-O-sulfotransferase 6
XP_011068417.1 dnaJ homolog subfamily C member 11
XP_011067624.1 UDP-NAG--dolichyl-phosphate N-acetylglucosaminephosphotransferase isoform X2
XP_011065542.1 prohormone-4
XP_011064646.1 alpha-2-macroglobulin receptor-associated protein
XP_011066277.1 uncharacterized protein C45G9.7
XP_011054141.1 stress-induced-phosphoprotein 1
XP_011063347.1 trypsin-1-like
XP_011063764.1 transmembrane protein 189
XP_011059987.1 zinc transporter ZIP9-A
XP_011061723.1 nitrilase and fragile histidine triad fusion protein NitFhit isoform X5
XP_011057182.1 dipeptidyl peptidase 3 isoform X2
XP_011065299.1 eukaryotic translation initiation factor 3 subunit L
XP_011060958.1 ATP-binding cassette sub-family F member 3 isoform X2
XP_011069277.1 LOW QUALITY PROTEIN: dihydropyrimidine dehydrogenase [NADP(+)]
XP_011061112.1 annexin B9
XP_011050995.1 39S ribosomal protein L28, mitochondrial isoform X2
XP_011052701.1 uncharacterized protein LOC105145068 isoform X2
XP_011065635.1 solute carrier family 2, facilitated glucose transporter member 1-like isoform X2
XP_011066832.1 PX domain-containing protein kinase-like protein isoform X2

Fig. S3

**Additional file 5: Table S2.** Pan-genomes across sequenced Holometabola.

<click for link to Table S2>

**Additional file 6: Table S3.** Analysis of *C. felis* proteins that did not cluster with other Holometabola.

<click for link to Table S3>

**Additional file 7: Table S4.** Elements of the *C. felis* microbiome and associated *Wolbachia* phylogeny estimation.

<click for link to Table S4>

**Additional file 8: Table S5.** Coverage of corrected PacBio reads against all 16,622 polished assembly contigs.

<click for link to Table S5>
